# BCell-Epitope-Prediction: A local, reproducible pipeline for multi-epitope vaccine design: integration and structural validation against a panel of PDB antigen-antibody complexes

**DOI:** 10.64898/2026.10.13.758921

**Authors:** Enzo Leonardo Edison Sierra Vera, Blanca Pueche Granados, Carmen Elena Gómez, Carlos Óscar S. Sorzano

## Abstract

Epitope-based subunit vaccine design requires chaining together heterogeneous analyses: B-cell antigenicity, topological accessibility, self-tolerance, allergenicity, T-helper/T-cytotoxic immunogenicity, and literature corroboration, historically scattered across separate, often web-only tools. This work presents a computational pipeline that integrates fourteen external tools across eight main phases, runs locally without network calls, and uses content-based self-caching. B-cell antigenicity is assessed via four engines (BepiPred-3.0, EpiDope, DiscoTope-3.0, and ScanNet), whose per-residue scores are fused into a single signal before thresholding.

Validation used a panel of seventeen real antigen-antibody complexes from the Protein Data Bank spanning five pathogenic species, with ground truth from structural contact and independent literature verification. Fusing the four engines’ scores, completed to a fixed candidate count, outperforms the best individual engine in F1 while reducing predicted residue volume. T-helper and T-cytotoxic immunogenicity are evaluated over each antigen’s complete sequence, independently of the B-cell candidates. The final construct retains a B-cell epitope in 13 of 17 structures, and a T-cell block in 13 of 17 antigens, of which 6 match literature-documented epitopes and 2 more show partial correspondence.

## 1. Introduction

Epitope-based subunit vaccine design (minimal peptide fragments capable of inducing a protective immune response, humoral or cellular) is an established alternative to attenuated or inactivated-pathogen vaccines, especially for agents with high biological risk, high mutability, or difficult recombinant production. This concentrates the immune response on regions with demonstrated or predicted protective value, avoids cross-reactivity against self-proteins, and reduces the experimental search space compared with phage display or large-scale ELISpot screening.

The immunoinformatics literature documents a well-known methodological bottleneck: a rigorous workflow must chain together B-cell antigenicity prediction (from sequence and structure, since conformational epitopes account for most real B-cell epitopes; Kumar et al. 2023), exclusion of antibody-inaccessible regions, discarding host-homologous candidates, allergenicity and N-glycosylation assessment, separate T-helper/T-cytotoxic immunogenicity assessment with processing evidence, and comparison against documented epitopes. Each step has reasonably good published tools, but no system organizes them in a reproducible, traceable way through to an assembled multi-epitope construct subjected to a safety check on the final sequence, something no per-peptide check can substitute for, since joining fragments can generate *de novo* motifs absent from any of them individually. Added to this is the usual dependence on web services or cloud APIs: availability bottlenecks, silent reproducibility loss when a remote service updates model weights, and a sequence-leakage risk for pathogens of biosafety relevance before publication.

EpiToolKit (Schubert et al. 2015), a web-based workbench published in this same journal, set the precedent for integrating multiple prediction tools into vaccine-design workflows, though it was built on tools available a decade ago. More recently, VaccineDesigner (Trygoniaris et al. 2025) integrates B-cell and T-cell (CTL/HTL) epitope prediction, multi-epitope assembly, population coverage, molecular mimicry screening, and proteasomal cleavage in a single web platform, with a scope close to the one presented here. To our knowledge, however, no open-access tool, including these two, combines the same screening layers with entirely local execution: VaccineDesigner and EpiToolKit are web services, that require sequence submission to a remote server, a limitation where sequence leakage before publication is a genuine biosafety concern. Nor does either validate epitope calls against real structural contact from resolved antigen-antibody complexes, relying instead on internal prediction metrics. Fully local execution and validation against physical structural evidence, not the breadth of screening layers, are the specific contribution of the pipeline presented here.

In this article, we propose a computational pipeline integrating fourteen external tools across eight main phases, run locally without network calls during inference. We assess whether fusing multiple B-cell antigenicity engines improves epitope detection over the best individual engine, quantify recovery of real B-cell epitopes over a panel of seventeen PDB complexes, verify T-cell immunogenicity against literature evidence, and document, with verified cause, the trade-offs and limitations found.

## 2. Materials and methods

The input type determines which antigenicity engines are activated in Phase 2 (Fig. 1). Phases 4b, 4c, 6, 6b, and 6c are informative (they do not filter candidates); the rest apply exclusion or threshold-based filtering. The self-tolerance re-check (Phase 4’s BLAST criterion, reapplied over the final binding window, sequence_f5) is common to both the HTL (Helper T Lymphocyte; Phase 5) and CTL (Cytotoxic T Lymphocyte; Phase 5b) pathways; population-coverage annotation applies only to CTL-pathway candidates, following NetCleave cross-checking (Phase 5b). Neither is a numbered phase; both are drawn as a single node in Fig. 1 where the two pathways converge, for layout compactness.

**Fig. 1.**
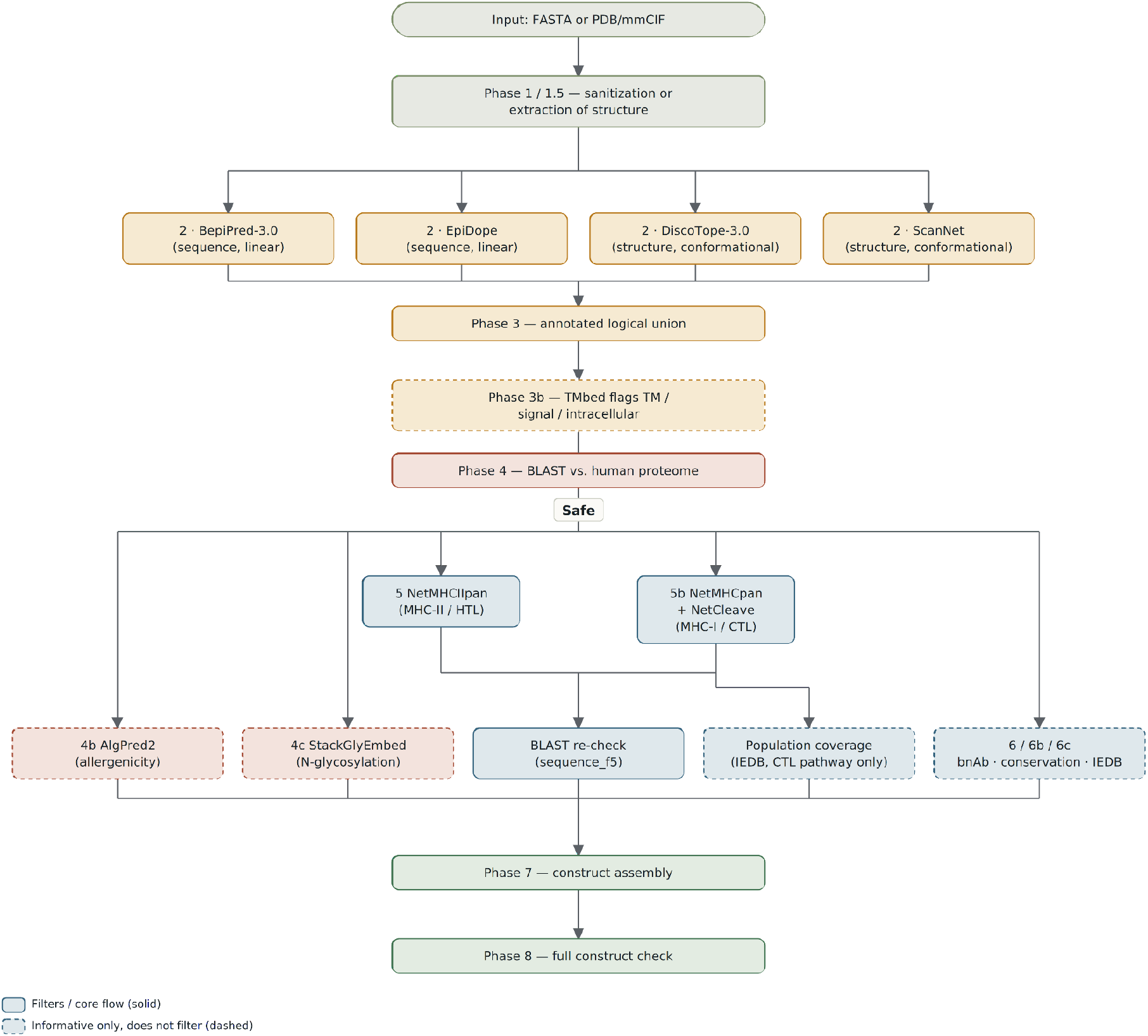
Complete pipeline flow (eight main phases, fourteen external tools).

Phase 1 (or 1.5, for a structure) sanitizes the input sequence or extracts the antigen’s resolved sequence and coordinates. Phase 2 scores every residue for B-cell antigenicity, using one to four engines depending on input type; Phase 3 fuses their normalized scores into non-redundant candidate regions. Phase 3b flags candidates overlapping a transmembrane, signal peptide, or intracellular segment as topologically inaccessible, without discarding them (informative only; Section 3.2). Phase 4 discards candidates with human-proteome homology; Phases 4b/4c annotate allergenicity and N-glycosylation risk. Phases 5/5b evaluate T-helper and T-cytotoxic immunogenicity over each antigen’s complete sequence, independently of which candidates survive Phase 3/4 (mirroring Phase 3b’s independence from the B-cell call), re-apply the self-tolerance check over the final binding window, and, for the T-cytotoxic pathway, verify antigen-processing evidence and estimate population coverage. Phases 6/6b/6c cross-check each B-cell candidate against published experimental evidence (broadly neutralizing antibodies, cross-strain conservation, and the IEDB). Phase 7 selects the best candidates per class, excludes any HTL/CTL candidate whose native range overlaps a higher-priority class already selected (cross-class deduplication), and assembles survivors into a single multi-epitope construct with full traceability; Phase 8 subjects it to a final, independent safety and quality check.

The system was implemented as a command-line orchestrator (pipeline.py, Python 3.10+; Fig. 1). Each computationally expensive phase self-caches via a content hash of its input, so an interruption never requires recomputing completed phases (e.g., loading three protein language models: ProteinBERT, Brandes et al. 2022; ESM-2 650M, Lin et al. 2023; ProtT5, Elnaggar et al. 2021). All external tools run as local subprocesses, with no network request during inference. A sequence FASTA activates the two sequence engines (BepiPred-3.0, EpiDope); a structure in structure_only mode activates the two structural engines (DiscoTope-3.0, ScanNet); a structure in default mode activates all four, additionally deriving an ATMSEQ sequence (the sequence actually resolved in the atomic coordinates, not the theoretical SEQRES) to feed the sequence engines; the same derived sequence Phases 3b, 5, and 5b scan in full. The entire validation panel used this third route. Complete technical parameters per phase (thresholds, BLAST E-values, window sizes, HLA allele panels and selection criteria) and each tool’s execution environment are detailed in Section S1, Supplementary Material. The full pipeline requires approximately 36 GB of disk space and completes a cold-cache, single-antigen run in under 8 minutes on a 16-core, GPU-free machine, independent of sequence length in this range (Supplementary Table S9).

B-cell antigenicity (Phases 2/3): four independent engines score every antigen residue. BepiPred-3.0 (Clifford et al. 2022; ESM-2 embeddings, Lin et al. 2023) and EpiDope (Collatz et al. 2021; ELMo embeddings, Peters et al. 2018) are sequence engines for linear epitopes; DiscoTope-3.0 (Høie et al. 2024; ESM-IF1 representation, Hsu et al. 2022) and ScanNet (Tubiana et al. 2022) are structural engines for conformational epitopes. Each engine’s raw score is min-max normalized (fixed global bounds, calibrated once over the full panel) and summed into a single fused signal, thresholded per engine combination by leave-one-out cross-validation (Section S1); adjacent calls are merged, with a minimum region length of 9 amino acids. Antigens with fewer than three regions above threshold are completed with the highest-scoring remaining 9-residue windows, up to three, plus up to three further windows kept in reserve for Phase 4 (evaluated in Section 3.1). Every candidate incorporating a structural signal is annotated with conformational_coverage_pct, the fraction of the real, possibly discontinuous, conformational patch captured within the synthesized window.

Biological safety filters (Phases 3b, 4, 4b/4c): Phase 3b runs TMbed (Bernhofer & Rost 2022) on the full sequence and flags candidates overlapping a transmembrane, signal-peptide, or intracellular segment, informatively (Section 3.2). Phase 4 subjects each region to a local BLASTp search (Camacho et al. 2009) against the human proteome (UniProt UP000005640; UniProt Consortium 2023); regions with maximum identity ≥75% are discarded for autoimmunity risk, the rest continue. Phases 4b (allergenicity, AlgPred2 hybrid mode; Sharma et al. 2021) and 4c (N-glycosylation, StackGlyEmbed; Nafi & Rahman 2025) are informative, annotating candidates without discarding on composition alone.

T-cell immunogenicity (Phases 5/5b): T-helper and T-cytotoxic evaluation is decoupled from the B-cell candidates of Phase 3/4; each antigen’s complete sequence is evaluated directly with NetMHCIIpan-4.3 (T-helper, panel of 27 HLA-DR/DRB3/DRB4/DRB5/DQ/DP alleles; Nilsson et al. 2023) and, in parallel, NetMHCpan-4.2 (T-cytotoxic, panel of 23 HLA-A/B/C alleles combining the 12 supertypes of Sidney et al. 2008, high-frequency HLA-C alleles from Rasmussen et al. 2014, and Weiskopf et al. 2013; Reynisson et al. 2020), since MHC presentation and antibody accessibility are biologically independent mechanisms (Section 3.2). A peptide is accepted as a “Valid Candidate” if it classifies as a strong or weak binder in at least 3 panel alleles. On the final binding window (sequence_f5, 15 aa in HTL, 8-11 aa in CTL), the Phase 4 self-tolerance filter is repeated, since a short risk motif can fail the 90% threshold at full-antigen scale yet be detected once recalculated over the shorter window; this second check does exclude candidates. Every CTL-pathway candidate is cross-checked with NetCleave (Amengual-Rigo & Guallar 2021) for proteasomal cleavage evidence, and annotated with population coverage via the IEDB methodology (Bui et al. 2006; Vita et al. 2019), weighting each allele by real frequency (Gonzalez-Galarza et al. 2020).

Literature corroboration and assembly (Phases 6/6b/6c, 7, 8): three informative mechanisms cross-check every "Safe" B-cell candidate against published evidence: broadly neutralizing HIV antibodies (LANL/CATNAP, Envelope antigens only), optional cross-strain conservation (a terminal coverage-cliff check automatically flags a non-native crystallization tail), and the IEDB’s B-cell bulk export (Vita et al. 2019). These are corroborative, not discovery-oriented: an empty result is expected for a novel target, not a phase failure. Phase 6 is restricted to HIV Envelope antigens (no LANL/CATNAP equivalent exists for other pathogens); Phase 6c’s IEDB cross-reference is broader (561 organisms, Section S1) but subject to the same limit. Phase 7 selects the 3 best candidates per class, excludes any HTL/CTL candidate whose range overlaps a higher-priority block already selected, and concatenates survivors with standard linkers (AAY intra-CTL, GPGPG intra-HTL and between classes, KK intra-B-cell; Livingston et al. 2002; Arai et al. 2001) in B-cell→HTL→CTL order, fully traceable to source. Phase 8 runs four further checks on the assembled sequence: allergenicity (AlgPred2; Sharma et al. 2021), toxicity (ToxinPred2; Sharma et al. 2022), intrinsic antigenicity (IApred; Miles et al. 2025), and signal-peptide absence (SignalP-6.0; Teufel et al. 2022). Two schemes were used for evaluation metrics: top-N recall (for initial exploratory benchmarking of each engine separately, N = real epitope size) and precision, recall, and F1 at the real production threshold (used throughout Section 3), calculated over predicted residues against the real epitope defined in the structural validation panel (Table 1). Both are aggregated in macro (per-antigen mean) and micro (true positives and residues summed panel-wide), since these can differ substantially when real epitope size is heterogeneous (8-27 residues here). For the fourteen antigens with at least one accepted T-cell candidate, it was verified whether the proposed candidate matches, at native position or literal sequence coincidence, a T-cell epitope already documented and validated in primary sources (IEDB, the LANL HIV CTL epitope database, articles identified via PMID). Internally predicted MHC binding affinity alone was not accepted as sufficient.

**Table 1.** Structural validation panel (17 PDB structures, 5 pathogenic species), with the primary publication describing each structure. Antigen abbreviations: AMA1, apical membrane antigen 1; CD4bs, CD4-binding site; CSP, circumsporozoite protein; fHbp, factor H binding protein; MPER, membrane-proximal external region; NHBA, neisserial heparin-binding antigen; RBD, receptor-binding domain. Each structure’s citation was verified against rcsb_primary_citation from the public RCSB PDB API and against PubMed (identical title, journal, and year in both sources). The sole exception is 8UP2 (fHbp site 2), deposited with no associated primary publication.

| PDB | Organism | Antigen | Antibody/Fab | Antigen residues | Primary citation (RCSB) | DOI |
| --- | --- | --- | --- | --- | --- | --- |
| 7CE2 | *C. tetani* | Tetanus toxin, fragment C | TT0067 | 446 | Wang *et al.* 2021 | 10.2210/pdb7CE2/pdb |
| 7OH1 | *C. tetani* | Tetanus toxin, LC-HN | TT110 | 845 | Pirazzini *et al.* 2021 | 10.2210/pdb7OH1/pdb |
| 3NGB | HIV-1 | *gp120*, *CD4bs* site | VRC01 | 349 | Zhou *et al.* 2010 | 10.2210/pdb3NGB/pdb |
| 4XAW | HIV-1 | *gp41*, *MPER* | 4E10 | 16 | Irimia *et al.* 2016 | 10.2210/pdb4XAW/pdb |
| 5O1R | *N. meningitidis* | NHBA | Fab 5H2 | 115 | Maritan *et al.* 2018 | 10.2210/pdb5O1R/pdb |
| 5O14 | *N. meningitidis* | fHbp, site 1 | Fab 1A12 | 229 | Lopez-Sagaseta *et al.* 2018 | 10.2210/pdb5O14/pdb |
| 8UP2 | *N. meningitidis* | fHbp, site 2 | JAR4 | 241 | No primary publication in RCSB ("to be published") | 10.2210/pdb8UP2/pdb |
| 2Q8A | *P. falciparum* | AMA1 (crystal form A) | 1F9 | 316 | Coley *et al.* 2007 | 10.2210/pdb2Q8A/pdb |
| 2Q8B | *P. falciparum* | AMA1 (crystal form B) | 1F9 | 292 | Coley *et al.* 2007 | 10.2210/pdb2Q8B/pdb |
| 9BJG | *P. falciparum* | AMA1 | 75B10 | 278 | Patel *et al.* 2025 | 10.2210/pdb9BJG/pdb |
| 9BJH | *P. falciparum* | AMA1 | 75C8 | 285 | Patel *et al.* 2025 | 10.2210/pdb9BJH/pdb |
| 8FDD | *P. falciparum* | CSP, NPNV repeat | Ky15.3 | 13 | Thai *et al.* 2023 | 10.2210/pdb8FDD/pdb |
| 7RXP | *P. falciparum* | CSP βCT | Fab1512 | 65 | Beutler *et al.* 2022 | 10.2210/pdb 7RXP/pdb |
| 6B5M | *P. falciparum* | CSP, junctional region | CIS43 | 13 | Kisalu *et al.* 2018 | 10.2210/pdb 6B5M/pdb |
| 7STR | SARS-CoV-2 | Nucleoprotein, site 1 | S24-1063 | 129 | Kim *et al.* 2024 | 10.2210/pdb 7STR/pdb |
| 7STS | SARS-CoV-2 | Nucleoprotein, site 2 | S24-1379 | 111 | Kim *et al.* 2024 | 10.2210/pdb 7STS/pdb |
| 7BEP | SARS-CoV-2 | Spike, RBD domain | COVOX-384 +S309 | 197 | Dejnirattisai *et al.* 2021 | 10.2210/pdb 7BEP/pdb |

## 3. Results

Structural validation panel and ground truth: the system’s ability to recover real B-cell epitopes was assessed over a panel of seventeen antigen-antibody complexes deposited in the Protein Data Bank (Berman et al. 2000) (Table 1), selected to span three broad, biologically distinct taxonomic categories (virus, bacterium, and a eukaryotic parasite). For each complex, the real epitope was defined automatically from the geometry of the deposited structure, with no intervention from the pipeline itself: a residue is considered part of the epitope if at least one of its heavy atoms is ≤4.0 Å from a heavy atom of the real binding partner (antibody Fab/Fv), identified via rules applied to the PDB file’s COMPND metadata and computed with Biopython (PDBParser + NeighborSearch; Cock et al. 2009) (Section S1 of the Supplementary Material details the complete procedure).

Structures were selected for taxonomic and epitope diversity rather than through an exhaustive search; full selection criteria are detailed in Supplementary Section S8.

### 3.1. Comparison of B-cell antigenicity engines against the score-fusion workflow

Before evaluating the pipeline, we tested Phase 3’s central design decision: fusing the four engines’ normalized scores into a single signal before thresholding, with the threshold calibrated by leave-one-out cross-validation. We evaluated each engine individually and the fused signal under the same criterion (real production threshold, before Phase 4) over the same set of ground-truth residues from the 17-structure panel (Table 2).

**Table 2.** Residue-level performance at the real production threshold, aggregated over the 17 structures. "Score fusion (Phase 3)" combines the four engines’ normalized scores by summation before thresholding, completing each antigen up to 3 candidate regions and up to 3 reserve regions when fewer than 3 cross the threshold (see Section 2); never a threshold or completion rule informed by prior knowledge of the true epitope’s size or location.

| Method | *Recall (macro)* | *Precision (macro)* | *F1 (macro)* | *Recall (micro)* | *Precision (micro)* | *F1 (micro)* |
| --- | --- | --- | --- | --- | --- | --- |
| *BepiPred -3.0* | 0.759 | 0.232 | 0.320 | 0.746 | 0.133 | 0.225 |
| *EpiDope* | 0.053 | 0.007 | 0.010 | 0.050 | 0.017 | 0.025 |
| *DiscoTope -3.0* | 0.303 | 0.209 | 0.236 | 0.326 | 0.259 | 0.289 |
| *ScanNet* | 0.097 | 0.142 | 0.112 | 0.122 | 0.233 | 0.160 |
| Score fusion (Phase 3) | 0.631 | 0.290 | 0.380 | 0.602 | 0.232 | 0.335 |

The fused signal outperforms every individual engine (each at its own published threshold) in F1, both micro (0.335 versus BepiPred-3.0’s 0.225) and macro (0.380 versus 0.320), though it does not win on raw recall: it calls 723 residues across the panel before Phase 4, versus BepiPred-3.0’s 1,566, trading recall for precision and F1 relative to BepiPred-3.0 (0.602 versus 0.746 micro recall; 0.232 versus 0.133 micro precision). Nor is there complete dominance by a single engine on every axis: BepiPred-3.0 has the best macro and micro recall, but DiscoTope-3.0 is close to the fused signal on micro precision (0.259 versus 0.232), concentrated in structures where DiscoTope-3.0 hits with high specificity (SARS-CoV-2 nucleoprotein, site 2, recall=0.875 and precision=0.583 with a single engine). EpiDope, despite being the only fully open-source engine among the two sequence-based ones, is the weakest in the panel by a wide margin, and its normalized score contributes little beyond noise to the fused signal. Additional combination rules (per-engine quality weighting, percentile-rank normalization, a pure OR (maximum) operator, and a per-fold logistic regression with freely learned weights as a theoretical ceiling) were tested exhaustively under the same leave-one-out scheme, and none exceeded the equal-weight sum reported here, indicating that the ceiling on this panel is set by the input signal’s own quality and redundancy (EpiDope near-noise, ScanNet’s dynamic range compressed in this panel), not by the combination rule itself. Removing EpiDope and refitting the fusion threshold from the remaining three engines does not improve the fused signal (full ablation in Supplementary Section S10); retaining it is an empirically neutral-to-favorable choice, not merely a reproducibility/licensing one. The F1 gain of score fusion over BepiPred-3.0 alone is statistically significant across the 17 paired antigens (Wilcoxon signed-rank test on per-antigen F1, W=17.0, p=0.026; bootstrap 95% CI of the mean per-antigen F1 difference, 20,000 resamples, seed=20260906: 0.020-0.106, excluding zero).

In *N. meningitidis* fHbp (site 1, PDB 5O14), BepiPred-3.0 reaches 0.333 (7 of 21) and DiscoTope-3.0 reaches 0.381 (8 of 21) recall on their own, each calling a different, non-overlapping half of the real epitope. The fused signal never sustains a 9-residue run above the calibrated threshold for this antigen, so the candidate set is completed with three top-up windows and three further reserve windows (Fig. 2; Section 2); three of these six windows do recover real epitope signal (156-164, 187-195, and 171-179, capturing 6, 5, and 2 of the 21 real residues respectively, 13 in total), but none of them is among the three highest-scoring, so Phase 7 selects a different combination for the assembled construct (Fig. 4).

**Fig. 2.**
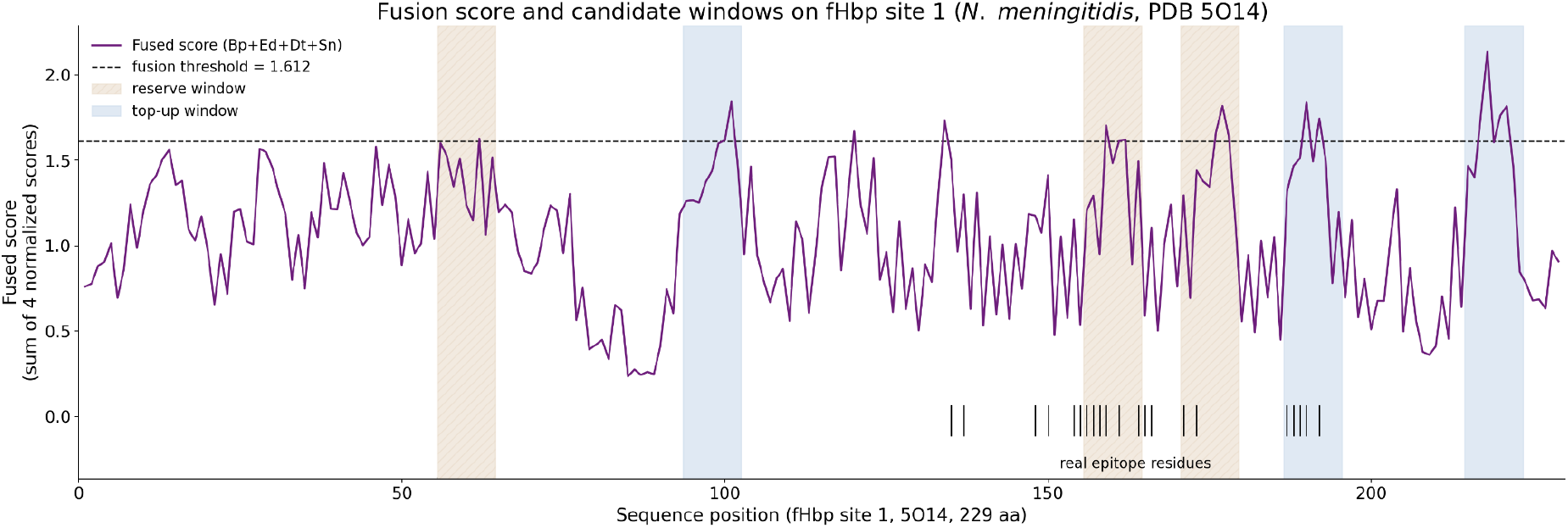
Fusion score and candidate windows on fHbp site 1 (*N. meningitidis*, PDB 5O14). The fused score (sum of the four normalized engine scores) crosses the calibrated threshold (dashed line, 1.612) only in short, scattered stretches, the longest just three residues (176-178), never sustaining the 9-residue minimum a region needs to qualify on its own. The candidate set is therefore completed with three top-up windows (highest-scoring non-overlapping 9-mers, blue) and three further windows kept in reserve (tan), shown against the 21 real epitope residues (tick marks, recomputed from structural contact). Three of the six candidates overlap the real epitope (156-164 and 187-195 recover 6 and 5 residues, respectively; 171-179 recovers 2, for 13 of 21 residues total); the other three (56-64, 94-102, 215-223) fall entirely outside it.

**Fig. 3.**
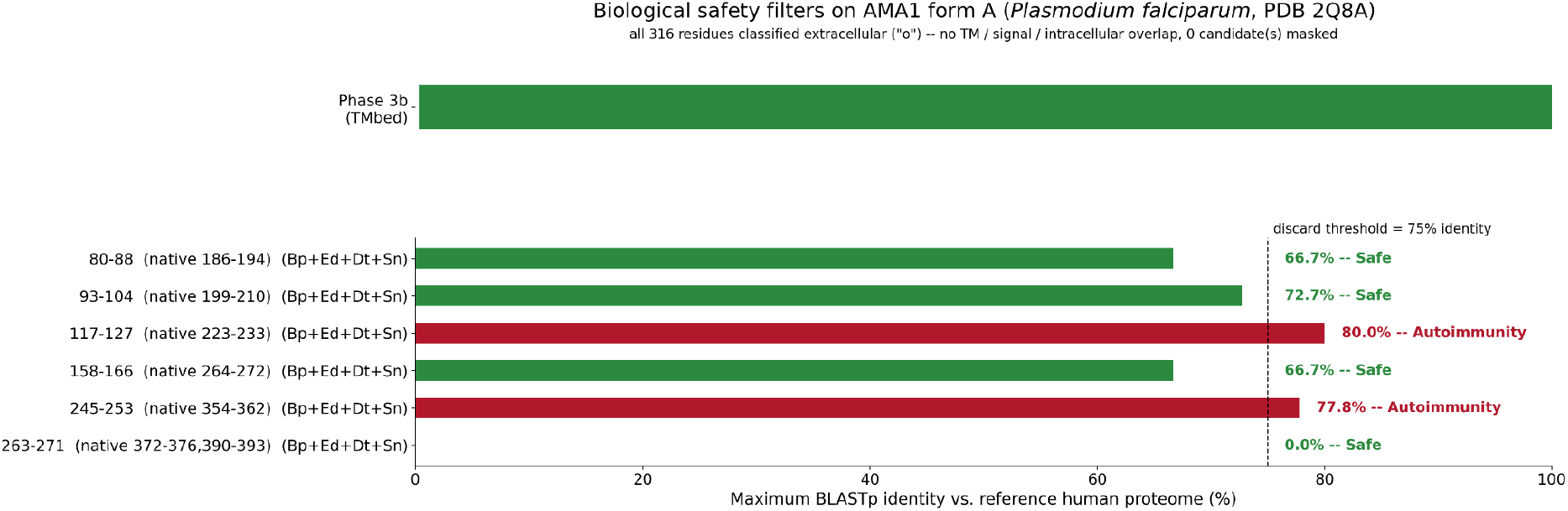
Biological safety filters on AMA1 form A (*Plasmodium falciparum*, PDB 2Q8A). Top: TMbed topology track over the full 316-residue antigen; all residues resolve extracellular, none masked. Bottom: maximum BLASTp identity against the reference human proteome for all six Phase 3 candidates, with the Phase 4 discard threshold (75%): four stay "Safe" (80-88, 93-104, 158-166 and 263-271, at 66.7%, 72.7%, 66.7% and 0.0% identity) and two are discarded as "Autoimmunity" (117-127 at 80.0%, 245-253 at 77.8%).

**Fig. 4.**
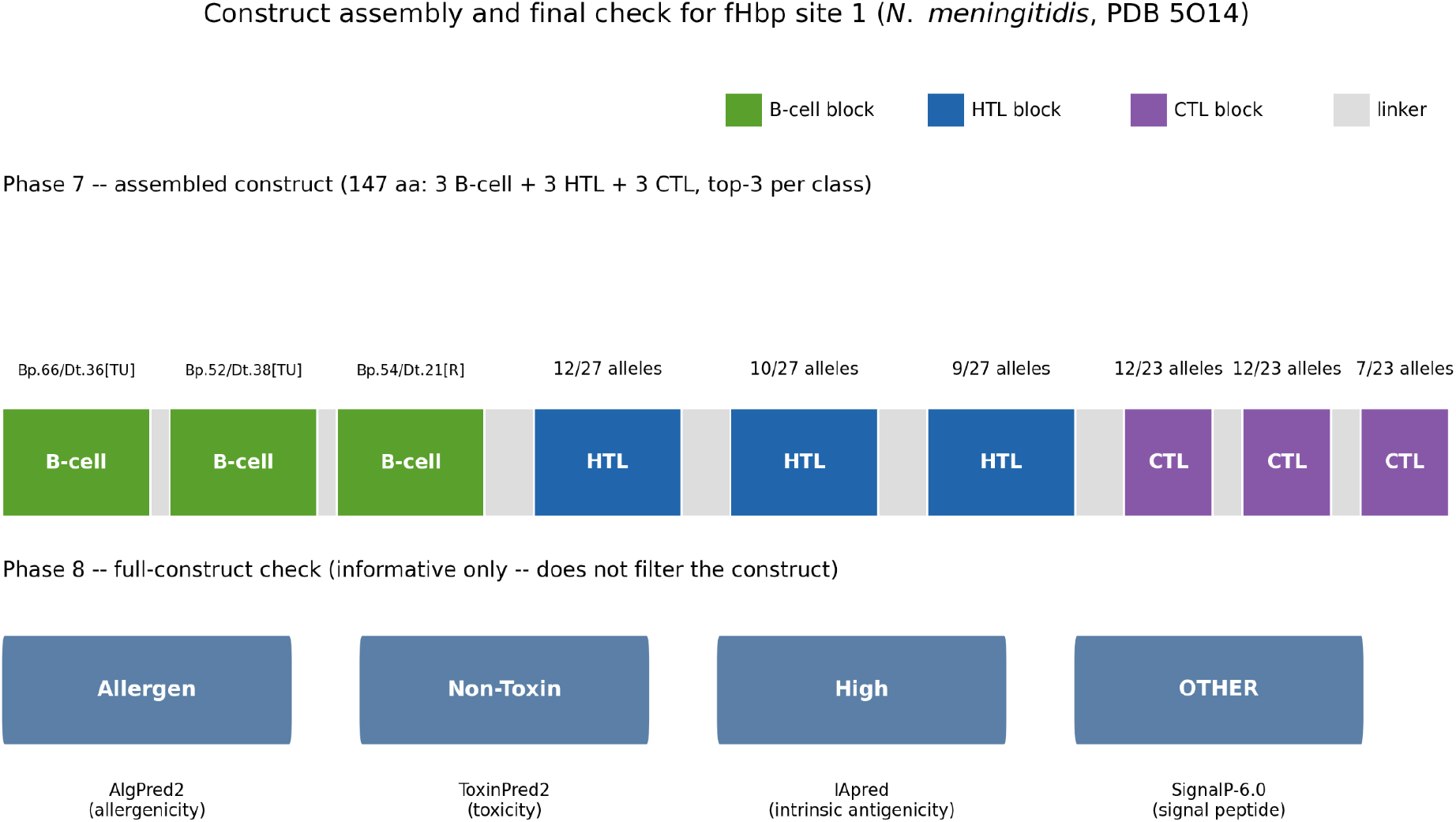
Construct assembly and final check for fHbp site 1 (*N. meningitidis*, PDB 5O14). Top: the assembled 147-aa construct (3 B-cell + 3 HTL + 3 CTL blocks + linkers, top-3 per class). All three B-cell blocks come from the completion mechanism of Fig. 2: two top-up windows (positions 212-226 and 184-198) and one reserve window (53-67), ranked and kept by fused score regardless of origin. Bottom: Phase 8’s four additional checks on the fully assembled sequence (Allergen, Non-Toxin, High intrinsic antigenicity, no signal peptide); all are informative only and do not filter the construct.

In tetanus toxin fragment C (7CE2), the fused signal reaches recall=0.714 (10 of 14 real residues), calling only 31 residues in total for the 14-residue true epitope; a high-precision result. Across the whole panel, this same threshold-and-completion logic brings total predicted volume to 723 residues (before Phase 4); three antigens (7OH1, 8UP2, 9BJG) end up with zero true positives despite having surviving candidates, but no antigen is left without at least one surviving candidate (Table 3); the direct, quantified cost of the precision achieved. This trade-off characterizes what score fusion buys the pipeline, and it is examined in full in Section 3.3.

**Table 3.**
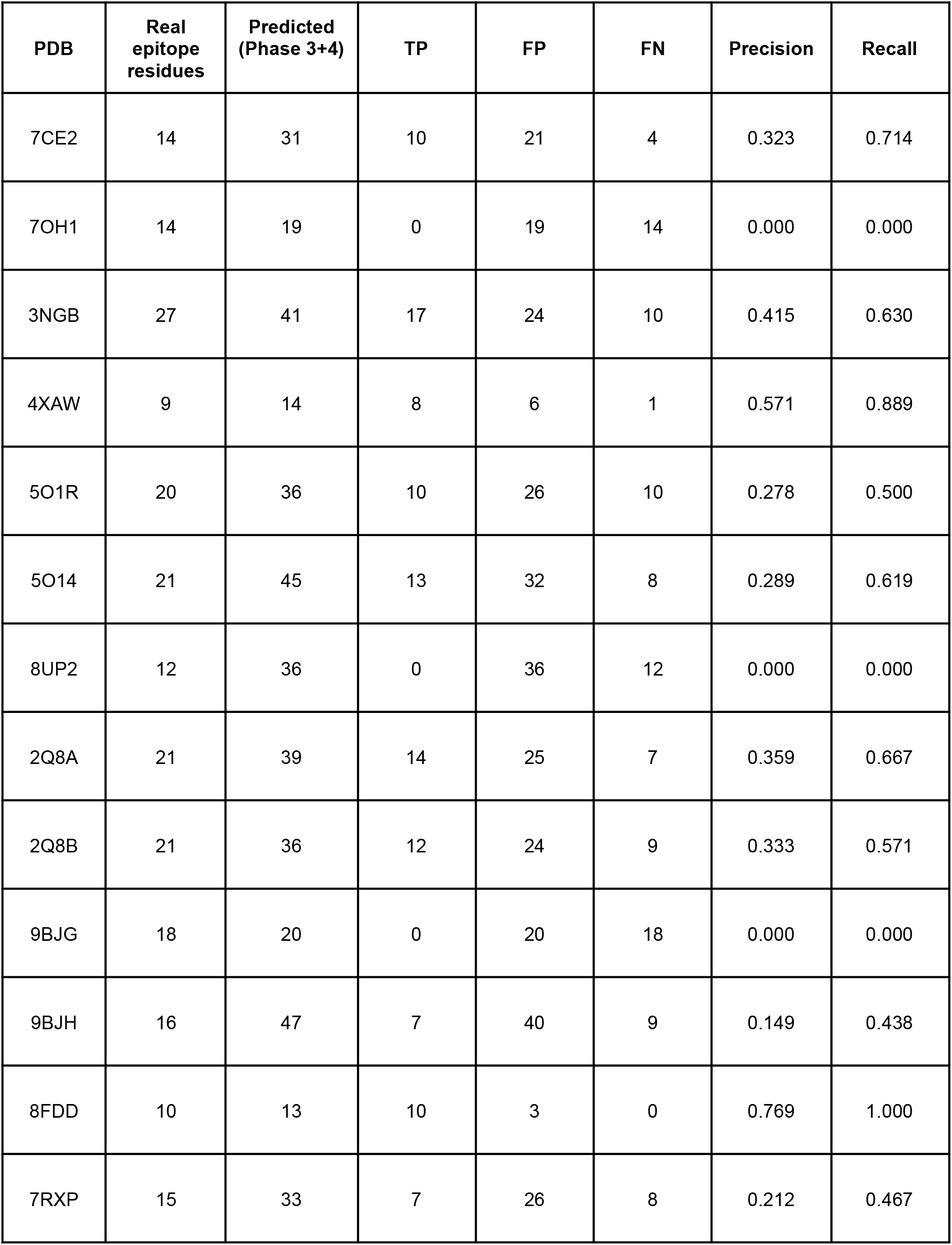
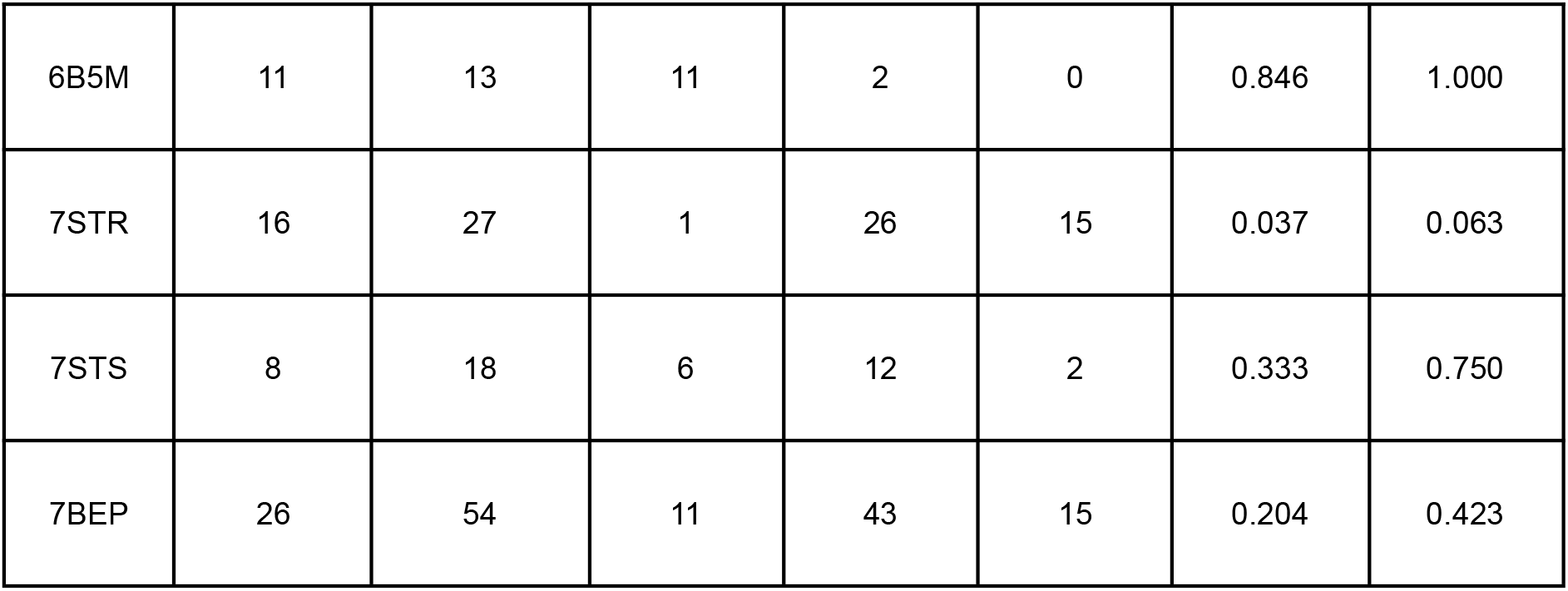
Residue-level agreement between the real structural epitope and the B-cell candidate set surviving Phase 3 (score fusion, completed up to 3 candidate regions plus up to 3 reserve regions per antigen) and Phase 4 (self-tolerance filter), by antigen, under the current production formula. TP, true positive; FP, false positive; FN, false negative (all residue counts, native PDB numbering). Precision = TP/(TP+FP); Recall = TP/(TP+FN).

### 3.2. Structural validation of the complete pipeline over the PDB panel

Across the 17-structure panel, every pipeline phase behaved as designed, with two findings that merit explanation.

First, in 6 of the 17 structures (tetanus toxin fragment C, gp41 MPER, both CSP fragments, and both SARS-CoV-2 nucleoprotein sites), TMbed labels 100% of candidates as intracellular despite these being surface/secreted antigens: its output alphabet has no class for "protein without topology," so a soluble protein resolves toward one of its two non-membrane labels (i/o). This does not discard any region or affect any panel result (Section S2).

Second, the self-tolerance filter (Phase 4) is far from cosmetic here: only 6 of 17 structures have 100% "Safe" candidates; the eleven exceptions lose one to four candidates each to human-proteome identity, in a 77.8-88.9% band (SARS-CoV-2 nucleoprotein site 2 loses four of six; CSP βCT and nucleoprotein site 1 lose three of six each) (Fig. 3).

To confirm this, we built an independent positive control outside the panel: a 20-aa fragment of M. tuberculosis GroEL2/Hsp65 with high similarity to its human ortholog HSPD1. It was correctly flagged as antigenic in Phase 3 and rejected in Phase 4 (89.47% identity, "Autoimmunity"). The panel’s 72 B-cell candidates show a continuous identity tail up to the threshold (48 of 72, 66.7%, in the 65-78% range), confirming Phase 4 operates over a populated region of identity space.

The assembled construct’s allergenicity check (Phase 8) flags Allergen in 16 of 17 final constructs (94.1%). Disaggregated against Phase 4b, 15 of those 16 already contained an individually-flagged B-cell block (propagating an existing value), consistent with a high background rate (32 of 42 B-cell blocks, 76.2%, flagged Allergen). Only one case (AMA1 form B, 2Q8B) is genuinely emergent: none of its three B-cell blocks was individually flagged, yet the assembled construct carries a real MERCI motif hit (score 0.5) absent from any individual block, direct evidence of the *de novo* junction-motif phenomenon motivating a construct-level check (Section 1). ToxinPred2 flags Toxin only in the two CSP-repeat constructs (8FDD, 6B5M; the same proline/asparagine-rich motif found in the licensed vaccines RTS,S/Mosquirix (RTS,S Clinical Trials Partnership 2015) and R21/Matrix-M (Datoo et al. 2021), both clinically safe), a composition-based false positive, not real toxicity.

The overlap between B-cell antigenicity and allergenicity is supported in the literature: VaxiJen (Doytchinova & Flower 2007) and AllerTOP (Dimitrov et al. 2013, 2014) apply the same auto/cross-covariance transform over the same descriptors, differing only in the target label; the distinction lies not in a structural property (Pomés 2010) but in Th2 polarization of the host response. This is not purely theoretical: clinical trials actively monitor IgE responses to licensed vaccine antigens (Aalberse et al. 2019; Pérez Chacón et al. 2024), underscoring why Phase 4b’s check is not a redundant formality.

As for T-cell immunogenicity, only 3 of 17 structures (17.6%) obtain no Phase 5/5b T-cell window at all: gp41 MPER (4XAW) and both CSP short fragments (8FDD, 6B5M). Since Phases 5/5b scan each antigen’s complete derived sequence independently of the B-cell call (Section 2), this is not a window-placement artifact: these three sequences are only 13-16 aa long, below the 15-aa HTL window and too short for meaningful MHC-I promiscuity regardless of core placement, an inherent floor of the crystallographic fragment (Section S4). Every other antigen yields at least one T-helper or T-cytotoxic block, including fHbp site 2 (8UP2, 241 aa) (Table S1); its comparatively weak MHC-I pathway is consistent with a systematic gap in fHbp’s T-cell record in IEDB (Section 3.3), not a search-strategy artifact.

### 3.3. Multi-epitope construct assembly and independent T-cell verification

At the individual-antigen level (Table 3), recall reaches or exceeds 0.7 in five of the seventeen structures (8FDD and 6B5M, both 1.0; 4XAW, 0.889; SARS-CoV-2 nucleoprotein site 2, 0.75; 7CE2, 0.714), each a short or single-epitope-dominated fragment where a 9-residue window covers a large share of the true epitope. Precision among the fourteen antigens with at least one true positive ranges from 0.037 (SARS-CoV-2 nucleoprotein site 1, 1 of 27 predicted) to 0.846 (6B5M, a 13-residue fragment where nearly the whole protein is antigenic); the remaining three have zero true positives, examined below.

Score fusion’s precision gain is real and concentrated in three antigens (4XAW, 8FDD, 6B5M), each exceeding 0.55 precision. The cost is concentrated, not spread evenly: three antigens (LC-HN/7OH1, fHbp site 2/8UP2, AMA1 75B10/9BJG) have zero true positives despite having surviving candidates; no antigen in the panel is left without one (Section 4). Construct-level retention (Phase 7’s top-3 selection per class) retains at least one real B-cell epitope residue in 13 of 17 structures (76.5%): full recall in the panel’s shortest, single-epitope-dominated fragments (4XAW, 8FDD, 6B5M), down to 0.125 in the weakest partial case (SARS-CoV-2 nucleoprotein site 1); the complete per-antigen breakdown is given in Supplementary Table S12. The four antigens with zero construct-level retention are LC-HN (7OH1), fHbp site 2 (8UP2), AMA1 75B10 (9BJG), and AMA1 75C8 (9BJH).

Three distinct root causes underlie these four zero-retention cases. In LC-HN (7OH1), no antigenicity engine generates a candidate region overlapping the real epitope: TT110 contacts two separate helices (597-607, 614-625) and an extending strand (Pirazzini et al. 2021), fragmenting the 14 contact residues into seven clusters of 1-4 residues each, none large enough to satisfy any engine’s 9-residue window support. 7OH1 is also the panel’s only cryo-EM structure (8.0 Å, versus 1.5-2.9 Å crystallographic for the rest), a caveat on the ground-truth definition’s precision here. In fHbp site 2 (8UP2) and AMA1 75B10 (9BJG), the completion mechanism generates a candidate overlapping the real epitope, but Phase 4 discards it on genuine human-proteome identity (77.8%): the real epitope region itself coincides with a site of real molecular mimicry with the host proteome, and admitting it would trade a real autoimmunity risk for two antigens’ worth of recall. We treat this as an accepted trade-off of the self-tolerance filter, calibrated panel-wide by design; 9BJG’s remaining margin is further limited by a real 5-residue fragmentation ceiling. In AMA1 75C8 (9BJH), two candidates with real overlap (capturing 4 and 3 of 16 real residues) survive Phase 4, but Phase 7’s ranking by fused score alone keeps three higher-scoring candidates with no overlap instead. The panel’s real IEDB cross-reference does return hits here, but for the three higher-scoring candidates Phase 7 already keeps, not for the two correct ones; an IEDB-based tie-breaker would not have rescued this case and could even reinforce the wrong choice, since no corroborating signal of any kind is available for the two correct candidates. This is classified as an evidence gap specific to this antigen, not a code defect. Score fusion’s residue-level advantage (Table 2) does not by itself raise construct-level retention: repeating Phases 3-4-7 with BepiPred-3.0 alone also retains 13 of 17 structures, though it recovers AMA1 75B10 (9BJG) where fusion does not, and vice versa for fHbp site 1 (5O14) (Supplementary Table S11).

Of the 14 antigens with at least one T-cell block, verification against primary sources produced heterogeneous results: solid confirmation in gp120 (a T-helper candidate nearly identical to the epitope documented in LANL since 1998), the four AMA1 structures (a T-helper candidate exactly overlapping the CD8 E6 epitope, validated by ELISpot/ICS; Sedegah et al. 2010, PMID 20735847), and SARS-CoV-2 nucleoprotein site 1 (documented HLA-B*07:02 epitopes). Nuanced results appear in Spike RBD (within the RBM region, but not the most-cited CD4 epitope) and LC-HN (a candidate overlapping the catalytic domain’s zinc-binding site, with CD4/CD8 epitopes characterized in the literature; Kerblat et al. 2000). A real absence was observed in tetanus toxin fragment C. fHbp site 1 is the panel’s only case with no T-cell literature available at all (confirmed via the public IEDB API; Vita et al. 2019): of 178 *N. meningitidis* entries, fHbp/GNA1870 has 13, all B-cell, none with an MHC allele, while other antigens from the same pathogen are characterized along both pathways (Fig. 5; Section S5).

**Fig. 5.**
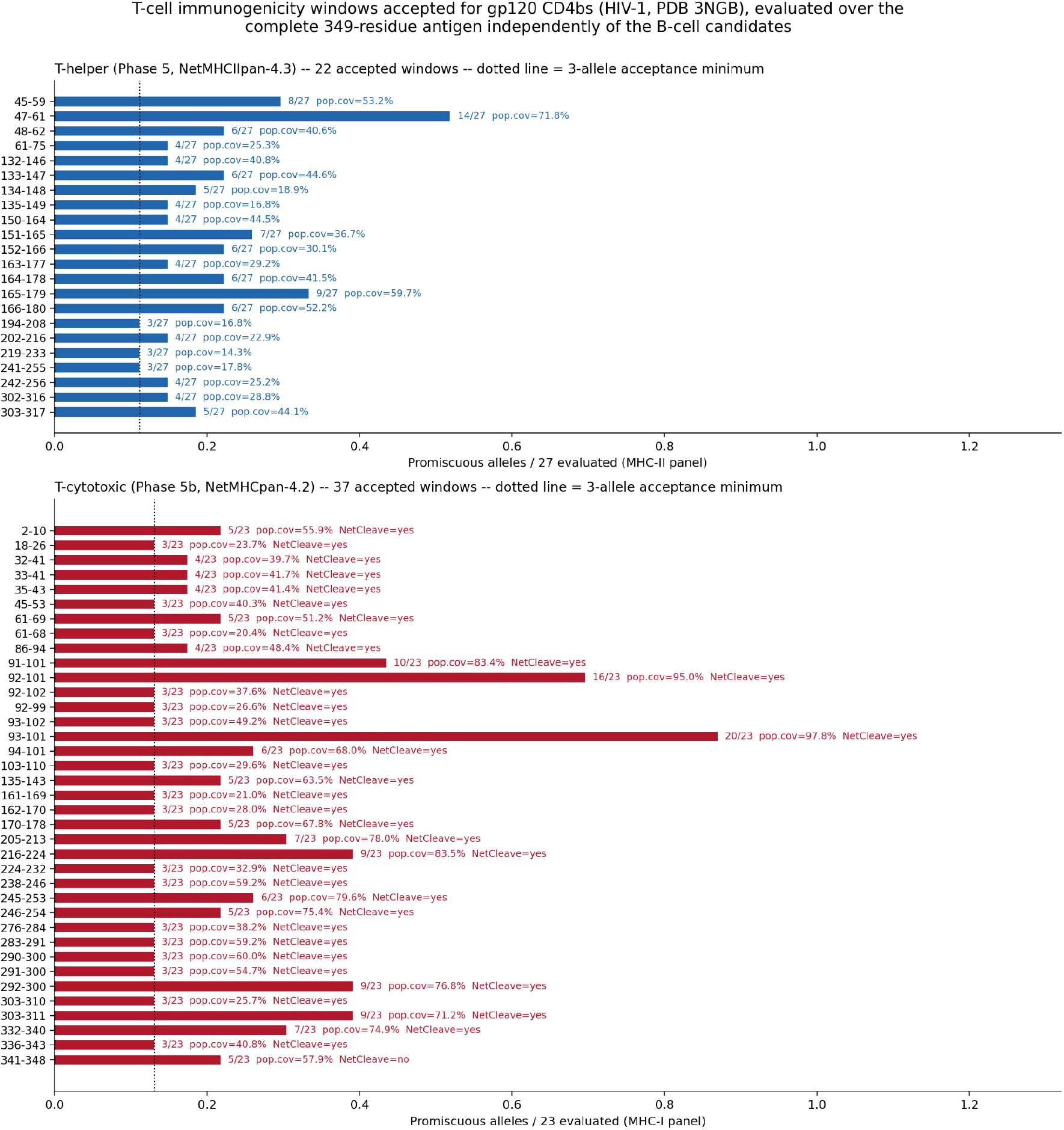
T-cell immunogenicity windows accepted for gp120 CD4bs (HIV-1, PDB 3NGB), evaluated over the complete 349-residue antigen independently of the B-cell candidates. Top: the 22 T-helper (Phase 5, NetMHCIIpan-4.3) windows accepted, promiscuity expressed as the fraction of the 27-allele reference panel bound as strong/weak binders; dotted line = 3-allele acceptance minimum. Bottom: the 37 T-cytotoxic (Phase 5b, NetMHCpan-4.2) windows over the 23-allele panel, annotated with population coverage and NetCleave C-terminal cleavage match. Phase 7 later selects only the top 3 candidates per class for the final construct.

## 4. Discussion

The results clarify where combining antigenicity engines adds value, and where it does not. Fusing the four engines yields the panel’s best F1, both macro and micro (Section 3.1), while BepiPred-3.0 alone reaches the highest raw recall and DiscoTope-3.0 the highest micro precision. Complementarity between a sequence and a structural engine appears when each independently detects a different, non-overlapping half of the same epitope (fHbp site 1, Fig. 2), a genuine multi-engine effect, not a thresholding artifact. Beyond the antigenicity call itself, value lies in screening layers no single engine provides: self-tolerance, allergenicity, N-glycosylation, T-helper/T-cytotoxic immunogenicity with processing evidence, and corroboration against documented epitopes, at the cost quantified in Sections 3.2-3.3. Of the panel’s four zero-retention antigens, only LC-HN (7OH1) reflects an absent signal; the other three reflect a downstream decision made after a correct candidate already exists (Section 3.3). A per-antigen adaptive threshold was rejected for Phase 2/3, for the same reason as ScanNet’s percentile threshold at residue level: it trades a fixed false-positive rate for one that varies unpredictably by antigen. The fixed, panel-wide threshold is an accepted trade-off: predictable behavior is kept at the cost of the antigen-specific misses characterized in Section 3.3. A narrow surviving candidate set after Phase 4 should signal a user to inspect discarded alternatives before trusting that antigen’s B-cell block.

Decoupling T-helper and T-cytotoxic evaluation from the B-cell candidates (Section 2) follows the same logic: anchoring MHC-II/MHC-I evaluation to score fusion’s short windows leaves little room for a 15-aa MHC-II window or meaningful MHC-I promiscuity in windows only 9-13 aa wide. Verified against literature-anchored ground truth: for AMA1, the documented CD8 E6 epitope (Sedegah et al. 2010) is recovered identically whether anchored to the B-cell window or run over the complete antigen, while for gp120 the full sequence recovers substantially more real signal (Section 3.3, Fig. 5). This mirrors Phase 3b’s independence from Phase 3: MHC presentation and antibody accessibility are independent mechanisms, with no basis for anchoring one to the other.

Construct assembly (Phase 7, Section 2) excludes, before top-3 selection, any HTL/CTL candidate whose native range overlaps a block already selected for a higher-priority class, verified to prevent duplication between AMA1’s 9BJG and 9BJH candidates. The completion and reserve mechanism (Section 2) guarantees every antigen at least one B-cell block in the final construct, even where Phase 4 discards most candidates for genuine homology (SARS-CoV-2 nucleoprotein site 2, 7STS, retains two safe candidates from six, one recovering 0.875 construct-level recall), but this guarantee does not extend to recovering the real epitope specifically, as the three zero-retention cases analyzed in Section 3.3 (8UP2, 9BJG, 9BJH) show: two are accepted self-tolerance trade-offs, the third a ranking miss with no corroborating signal available for either correct candidate.

In the panel’s two shortest antigens, CSP βCT (7RXP, 65 aa) and NHBA (5O1R, 115 aa), Phase 7’s cross-class deduplication removes most HTL/CTL candidates: 7RXP’s construct has three B-cell blocks and none from HTL/CTL, and 5O1R’s has three B-cell blocks against one HTL and three CTL blocks (Table S2). This is an accepted trade-off: the completion mechanism supplies B-cell candidates covering a large share of these short antigens’ sequence, and the B-cell→HTL→CTL priority order keeps them over T-cell windows selected purely by allele promiscuity. A class-aware allocation rule reserving minimum HTL/CTL slots is a possible extension, not pursued here, since choosing which class to sacrifice under a fixed construct length has no principled answer without ground truth. The cross-strain conservation mechanism (Phase 6b), demonstrated on the panel’s two HIV-1 structures over 2,994 Env sequences from LANL/CATNAP (Section S6), automatically distinguishes a non-native flanking sequence from the real epitope’s own variability. 4XAW’s crystallized peptide carries non-native flanking residues beyond the native gp41 MPER motif (671-682); its BLASTp hits show a clean coverage cliff, near-total alignment across the native core collapsing to near-zero over the flanking residues. Evaluating conservation over the full crystallized peptide yields 0.23% breadth, an artifact of the flanking residues; the detector flags this directly from the BLASTp hits and reports conservation over the supported native sub-range alone, 60.1%, consistent with MPER’s status as a target of broadly neutralizing antibodies (4E10, 2F5, 10E8). A manual check of the PDB deposition header was tested and rejected: ENGINEERED appears in 15 of 17 structures without discriminating anything, and SYNTHETIC produces a real false positive on the CSP NPNV repeat (8FDD), chemically synthesized but 100% native in sequence.

These limitations share a common thread: each was found because this work validated against evidence external to the pipeline’s own criteria. Real structural contact, real literature-documented epitopes, a real conservation panel, a real PDB deposition header; rather than trusting any phase’s internal consistency. External validation is indispensable not only for evaluating the system as a whole, but for finding which phase needs scrutiny, and recognizing when a limitation is genuinely resolved rather than merely moved.

## 5. Conclusions

The eight-main-phase architecture was implemented and run over the validation panel’s 17 structures, integrating fourteen external tools locally, traceably, and with self-caching, with no network call during inference. Combining antigenicity engines improves the sensitivity-specificity trade-off of B-cell epitope detection relative to every individual engine, in both micro and macro F1, though it still predicts several-fold more residues than are truly antigenic in the panel’s most diluted cases (Table 3), and BepiPred-3.0 alone reaches the highest raw recall. We identified and characterized the mechanism responsible: complementary, non-redundant rescue in a subset of antigens with mixed linear/conformational epitopes.

The validation scheme, with ground truth derived from real physical contact in 17 PDB antigen-antibody complexes spanning five phylogenetically distinct pathogenic species, enabled quantification of the system’s B-cell epitope recovery capacity. Multi-epitope construct assembly, with full traceability of every segment back to its origin, retains a real B-cell epitope in 13 of the panel’s 17 structures (76.5%). T-helper and T-cytotoxic immunogenicity, evaluated independently over each antigen’s complete sequence and verified against literature-anchored ground truth, is confirmed in 14 of 17 antigens. The assembled construct’s allergenicity check showed that nearly all of its alarm verdicts are explainable through propagation of an already-reported per-peptide value; a single case is emergent at the construct level, and it carries a real, documented IgE motif absent from every individual block, direct evidence of the *de novo* junction-motif risk this construct-level check exists to catch.

Cross-class duplication in construct assembly is resolved by an explicit deduplication rule, verified against the case that motivated it. Phase 7’s ranking by fused score alone leaves two different residual patterns in the affected antigens. In fHbp site 2 (8UP2) and AMA1 75B10 (9BJG), the only candidate overlapping the real epitope carries genuine identity to a human protein, and Phase 4 discards it as designed; we treat this as an accepted trade-off of prioritizing autoimmunity safety over recall in these two antigens. In AMA1 75C8 (9BJH), two overlapping candidates survive Phase 4, but Phase 7 keeps three higher-scoring alternatives instead; the panel’s real IEDB cross-reference corroborates those higher-scoring alternatives, not the correct candidates, so an evidence-aware tie-breaker remains a concrete target only where corroborating evidence actually favors the real epitope, which is not the case here. A real HIV-1 Env conservation panel, demonstrated for both HIV-1 structures in the panel, additionally surfaced a documentation caveat specific to engineered crystallization constructs, resolved by detecting the non-native flanking sequence automatically from the BLASTp hits’ own coverage pattern, without relying on PDB deposition metadata. Taken together, these results support the conclusion that the developed system meets its objective: a reproducible, locally run tool, transparent about its own limitations, and validated with evidence that the candidates it proposes correspond to biologically real epitopes, not artifacts of the prediction process itself. Its modular architecture and its validation scheme are extensible to antigen panels and pathogen families beyond this panel.

## Data Availability

The pipeline’s source code is freely available for non-commercial academic use at https://github.com/Lvera-code/BCell-Epitope-Prediction under a CC BY-NC 4.0 license, archived at Zenodo (https://doi.org/10.5281/zenodo.22549053, commit c41690b, tag v1.0-submission). The results reported in Tables 2 and 3 correspond to this tagged repository state. The seventeen PDB structures used for structural validation are publicly available at the Protein Data Bank under accession codes 7CE2, 7OH1, 3NGB, 4XAW, 5O1R, 5O14, 8UP2, 2Q8A, 2Q8B, 9BJG, 9BJH, 8FDD, 7RXP, 6B5M, 7STR, 7STS, and 7BEP, and are additionally cited individually in the reference list (DataCite format) with their respective DOIs.

