## Supplementary Material for "BCell-Epitope-Prediction: A local, reproducible pipeline for multi-epitope vaccine design: integration and structural validation against a panel of PDB antigen-antibody complexes"

### S1. Technical parameters by phase.

Execution environments. The fourteen external tools were run in their own isolated environment (dedicated venv or conda), given the incompatible range of ages across their dependency stacks (from Python 3.6 with TensorFlow 1.14 to modern PyTorch 2.x), which ruled out a single shared environment.

Phases 2/3; B-cell antigenicity: sliding window of 9 amino acids, with tolerance of up to 2 residues below threshold within the window. Decision thresholds: BepiPred-3.0, 0.1512; EpiDope, 0.818; DiscoTope-3.0 (calibrated score), 0.90; ScanNet, adaptive threshold by 90th percentile (default), computed per antigen since ScanNet does not publish an absolute reference threshold. Any resulting region shorter than 9 amino acids is discarded before continuing. Contributing-engine annotation: Bp=BepiPred-3.0, Ed=EpiDope, Dt=DiscoTope-3.0, Sn=ScanNet; combinations joined by + (e.g., "Bp+Ed+Dt+Sn" when all four contribute). If fewer than 3 regions cross the threshold for a given antigen, the candidate set is completed up to 3 with the highest-scoring non-overlapping 9-amino-acid windows elsewhere in the sequence (candidate_source = top_up), and up to 3 further non-overlapping windows are kept in reserve (candidate_source = reserve) for Phase 4 to draw on if a higher-ranked candidate is discarded there; both use the same greedy selection by mean fused score, with no separate threshold of their own. The completion target (3) reuses the same per-class limit already applied in Phase 7 (Settings.CONSTRUCT_TOP_N_PER_CLASS), not an independent parameter.

Phase 3b; Membrane topology: TMbed (Bernhofer & Rost 2022, ProtT5-XL-U50 encoder) runs on the antigen's full sequence, not on each candidate peptide separately. Output alphabet: 5 classes per residue (B/b, H/h for transmembrane helix or strand, S for signal peptide, i/o for non-transmembrane inside/outside). There is no sixth class for "protein without topology."

Phase 4; Self-tolerance: dynamic selection of BLASTp algorithm and E-value by peptide length: ≤30 amino acids with blastp-short and a lenient E-value (50); 31-100 amino acids with standard blastp and E-value 0.1; >100 amino acids with standard blastp and E-value 0.05. A hit is only counted toward the maximum identity percentage if its alignment covers at least 90% of the length of the queried peptide. Discard threshold: maximum identity ≥75% against the reference human proteome (UniProt, canonical proteome UP000005640).

Phase 4b; Allergenicity: AlgPred 2.0, hybrid mode by default. The Hybrid Score combines a machine-learning component (Random Forest on amino acid composition), a homology component (local BLASTp, E-value $1\times10^{-6}$ against a database of allergens with documented IgE reactivity), and a motif component (MERCI search for documented IgE motifs; Vens, Rosso & Danchin 2011), with a decision threshold of 0.3. The machine-learning mode was discarded as a standalone criterion after confirming that it produces "Allergen" verdicts unsupported by real homology or motif.

Phase 4c; N-glycosylation: scan of the canonical sequon Asn-X-[Ser/Thr] (X≠Proline, including overlapping sequons). Each candidate site is evaluated with StackGlyEmbed (a cascade classifier over combined ProteinBERT, ESM-2 650M, and ProtT5 representations).

Phase 5; T-helper (MHC-II): NetMHCIIpan-4.3, panel of 27 HLA-DR/DRB3/DRB4/DRB5/DQ/DP alleles, selected for population representativeness according to the IEDB reference panel. Length-based routing: ≤40 amino acids in exact-peptide mode; >40 amino acids in protein mode (15-amino-acid sliding window). The 40-residue cutoff was set after confirming that peptide mode causes a binary buffer overflow for inputs >55 amino acids with this panel. Alleles flagged Inverted by NetMHCIIpan itself are excluded. Acceptance threshold: strong or weak binder (default %Rank thresholds) in at least 3 alleles of the panel in normal orientation.

Phase 5b; T-cytotoxic (MHC-I): NetMHCpan-4.2, panel of 23 HLA-A/B/C alleles: 12 class I supertypes from Sidney et al. (2008) + 11 high population-frequency HLA-C alleles from Rasmussen et al. (2014), with a population frequency criterion ≥1% from IEDB, complemented with Weiskopf et al. (2013) for HLA-C representation. %Rank thresholds: 0.5 strong / 2.0 weak. Additional cross-check with NetCleave (model pre-trained on HLA mass spectrometry data, IEDB/UniProt/UniParc) to verify proteasomal cleavage at the residue following the binding core. Informative annotation does not condition the acceptance verdict.

Deduplication and self-tolerance re-check (Phases 5/5b). Protein-mode windows are deduplicated by grouping on the exact 9-amino-acid binding core and exact promiscuity, keeping the row with the best (lowest) %Rank in each group. On the sequence_f5 column (the final window: 15 aa in HTL, 8-11 aa in CTL), blast_engine.filter_self_tolerant is run, a second self-tolerance filter with the same blast_db/identity_threshold as Phase 4. This is necessary because Phase 4 operates on the Phase 3 candidate region (approximately 20-70 amino acids): a short risk motif buried within that longer region can have an alignment coverage so low at that scale (13-55% of the original length) that it never crosses the 90% threshold required in Phase 4. It is only detected by recalculating coverage over the length of sequence_f5. This second check does exclude candidates. Population coverage is estimated, after the previous filter, following the Population Coverage methodology of the IEDB (Immune Epitope Database) (Bui et al. 2006), weighting each allele by its real frequency in the Allele Frequency Net Database. Panel alleles without an available frequency in the source are excluded from the calculation and are never assumed to be 0.

Phases 6/6b/6c; Corroboration: Phase 6 (HIV bnAb): substring match, minimum 6-amino-acid overlap, against linear epitopes from the LANL HIV Molecular Immunology database; appends IC50/breadth from CATNAP. Phase 6b (conservation, optional): identity ≥90% against a user-supplied multi-FASTA panel of variants of the same pathogen; restricted to B-cell candidates, a terminal coverage-cliff check flags a candidate whose BLASTp hits (at the same identity threshold, before the query-coverage filter) show a contiguous run of 2 to 5 residues at one end with per-position coverage below 10% while the rest of the candidate stays above 80%, a signature of a non-native crystallization or cloning tail; when flagged, conservation is recomputed over the supported sub-range alone and both figures are reported. Phase 6c (IEDB): same 6-amino-acid overlap criterion against a locally filtered subset of the IEDB's B-cell bulk export (3,429 records, 561 organisms), restricted to records with a neutralization or protection response measure and a positive qualitative outcome.

Phase 7; Assembly: selection of the 3 best candidates from each class (B-cell, HTL, CTL). Linkers: AAY intra-CTL, GPGPG intra-HTL and between classes (Livingston et al. 2002), KK intra-B-cell. Block order: B-cell → HTL → CTL. No adjuvant, by active design decision. The function optionally supports a rigid EAAAK linker (Arai et al. 2001) to incorporate an adjuvant in future iterations.

Phase 8; Construct check: AlgPred 2.0 (reused from Phase 4b); ToxinPred2 (Raghava group, threshold 0.6, recommended by its authors for full-length proteins); IApred (Miles et al. 2025, replacing VaxiJen since the latter is not open-source or available as a local version); SignalP-6.0 (slow_sequential mode).

Ground-truth definition. An antigen residue is considered part of the real epitope if at least one of its heavy (non-hydrogen) atoms is ≤4.0 Å from a heavy atom of the real binding partner's chain or chains, computed with Biopython (PDBParser + NeighborSearch). Identification of the real binding partner via two rules applied to COMPND metadata:

1. Exclusion of chains whose molecular name is identical to that of the reference antigen (crystal-packing contact, not an antibody).

2. Regrouping of remaining chains by exact molecular name, keeping from each group the copy with the highest number of real contact residues, requiring a minimum of 2 contact residues to accept a chain as a real binding partner.

### S2. Phase 3b — Verification of topology masking.

In 6 of the panel's 17 structures (tetanus toxin fragment C, gp41 MPER, the CSP NPNV repeat and junctional region, and both sites of the SARS-CoV-2 nucleoprotein), TMbed labels 100% of that protein's candidate regions as intracellular. Verified directly: the number of masked regions is identical to the number of Phase 3 candidate regions in these 6 cases. The annotation never discards any real region in this specific panel; nor in any other, since Phase 3b is informative only by design and never excludes candidates automatically (Section 3.2 of the main body). Every residue that is neither transmembrane nor a signal peptide is forced, by the classifier's own design, to resolve as *i* (inside) or *o* (outside); never as "none." For a soluble, secreted, or extracellular protein with no real transmembrane segment, the model lacks a real topological anchor from which to decide that binary disjunction residue by residue, and in this panel it consistently resolves toward *i* in the 6 affected proteins. This is not a poorly defined "default" class in the code, but a consequence of the model's own output space lacking an alternative.

### S3. Complete per-antigen tables.

| Antigen (PDB) | Phase 4: safe/ candidates (%) | Phase 5 HTL promiscuous (mean cov.) | Phase 5 best candidate (cov.) | Phase 5b CTL promiscuous (mean cov.) | Phase 5b % NetCleave match |
|---|---|---|---|---|---|
| Tetanus toxin fragment C (7CE2) | 3/3 (100.0%) | 93 (27.7%) | 64.8% | 106 (54.2%) | 99.1% |
| Tetanus toxin LC-HN (7OH1) | 2/3 (66.7%)* | 222 (30.2%) | 81.6% | 198 (50.9%) | 99.5% |
| gp120 CD4bs, HIV-1 (3NGB) | 4/6 (66.7%)* | 22 (35.3%) | 71.8% | 37 (54.0%) | 97.3% |
| gp41 MPER, HIV-1 (4XAW) | 1/1 (100.0%) | 0 (-) | - | 0 (-) | - |
| NHBA (5O1R) | 4/6 (66.7%)* | 18 (24.5%) | 52.0% | 16 (43.0%) | 100.0% |
| fHbp site 1 (5O14) | 5/6 (83.3%)* | 53 (30.9%) | 75.3% | 29 (52.4%) | 100.0% |
| fHbp site 2 (8UP2) | 4/6 (66.7%)* | 56 (31.1%) | 75.3% | 30 (46.4%) | 100.0% |
| AMA1 1F9 form A (2Q8A) | 4/6 (66.7%)* | 39 (32.0%) | 66.0% | 34 (49.2%) | 100.0% |
| AMA1 1F9 form B (2Q8B) | 3/3 (100.0%) | 35 (31.2%) | 66.0% | 32 (50.9%) | 100.0% |
| AMA1 75B10 (9BJG) | 2/3 (66.7%)* | 32 (32.6%) | 66.0% | 24 (53.7%) | 100.0% |
| AMA1 75C8 (9BJH) | 5/6 (83.3%)* | 36 (33.0%) | 66.0% | 27 (51.7%) | 100.0% |
| CSP NPNV repeat (8FDD) | 1/1 (100.0%) | 0 (-) | - | 0 (-) | - |
| CSP βCT (7RXP) | 3/6 (50.0%)* | 8 (21.8%) | 31.7% | 2 (58.8%) | 100.0% |
| CSP junctional region (6B5M) | 1/1 (100.0%) | 0 (-) | - | 0 (-) | - |

| | | | | | |
|---|---|---|---|---|---|
| SARS-CoV-2 nucleoprotein, site 1 (7STR) | 3/6 (50.0%)* | 14 (40.7%) | 71.9% | 10 (51.8%) | 100.0% |
| SARS-CoV-2 nucleoprotein, site 2 (7STS) | 2/6 (33.3%)* | 11 (41.4%) | 71.9% | 17 (51.0%) | 100.0% |
| Spike RBD, SARS-CoV-2 (7BEP) | 3/3 (100.0%) | 39 (25.5%) | 49.6% | 33 (50.4%) | 100.0% |

Table S1. Self-tolerance (Phase 4) and T-helper/T-cytotoxic immunogenicity (Phases 5/5b) by antigen. 17-structure validation panel. T-helper and T-cytotoxic evaluation runs over each antigen's complete sequence, independently of the Phase 3/4 B-cell candidates (Section 2). *Antigens where at least one Phase 3 candidate was discarded by Phase 4 (≥75% identity against the reference human proteome; Section S1).

| Antigen (PDB) | Construct length (aa) | B/HTL/CTL blocks | AlgPred2 (Phase 8) | ToxinPred2 | IApred | SignalP-6.0 |
|---|---|---|---|---|---|---|
| Tetanus toxin fragment C (7CE2) | 151 | 3/3/3 | Allergen (no MERCI/BLAST) | Non-Toxin | High | OTHER |
| Tetanus toxin LC-HN (7OH1) | 131 | 2/3/3 | Allergen (BLAST) | Non-Toxin | High | OTHER |
| gp120 CD4bs, HIV-1 (3NGB) | 155 | 3/3/3 | Allergen (MERCI) | Non-Toxin | Moderate | OTHER |
| gp41 MPER, HIV-1 (4XAW) | 16 | 1/0/0 | Non-Allergen | Non-Toxin | Not evaluated (<20 aa) | OTHER |
| NHBA (5O1R) | 107 | 3/1/3 | Allergen (no MERCI/BLAST) | Non-Toxin | High | OTHER |
| fHbp site 1 (5O14) | 147 | 3/3/3 | Allergen (no MERCI/BLAST) | Non-Toxin | High | OTHER |
| fHbp site 2 (8UP2) | 147 | 3/3/3 | Allergen (no MERCI/BLAST) | Non-Toxin | High | OTHER |
| AMA1 1F9 form A (2Q8A) | 150 | 3/3/3 | Allergen (MERCI) | Non-Toxin | High | OTHER |

| | | | | | | |
|---|---|---|---|---|---|---|
| AMA1 1F9 form B (2Q8B) | 156 | 3/3/3 | Allergen (MERCI) | Non-Toxin | High | OTHER |
| AMA1 75B10 (9BJG) | 132 | 2/3/3 | Allergen (MERCI) | Non-Toxin | High | OTHER |
| AMA1 75C8 (9BJH) | 149 | 3/3/3 | Allergen (MERCI) | Non-Toxin | High | OTHER |
| CSP NPNV repeat (8FDD) | 13 | 1/0/0 | Allergen (no MERCI/BLAST) | Toxin | Not evaluated (<20 aa) | OTHER |
| CSP βCT (7RXP) | 47 | 3/0/0 | Allergen (no MERCI/BLAST) | Non-Toxin | Moderate | OTHER |
| CSP junctional region (6B5M) | 13 | 1/0/0 | Allergen (no MERCI/BLAST) | Toxin | Not evaluated (<20 aa) | OTHER |
| SARS-CoV-2 nucleoprotein, site 1 (7STR) | 146 | 3/3/3 | Allergen (MERCI) | Non-Toxin | High | OTHER |
| SARS-CoV-2 nucleoprotein, site 2 (7STS) | 106 | 2/3/1 | Allergen (MERCI) | Non-Toxin | High | OTHER |
| Spike RBD, SARS-CoV-2 (7BEP) | 152 | 3/3/3 | Allergen (no MERCI/BLAST) | Non-Toxin | High | OTHER |

Table S2. Construct assembly (Phase 7) and final check (allergenicity, toxicity, intrinsic antigenicity, and signal peptide (Phase 8)) by antigen. 17-structure validation panel. Allergen (no MERCI/BLAST) cases are those where the result rests on the ML/composition classifier, discussed as a composition-based false-positive pattern in Section 3.2 of the main body. B/HTL/CTL block counts below 3 per class reflect either an antigen too short to generate three independent candidates (4XAW, 8FDD, 6B5M) or Phase 7's cross-class deduplication rule, which excludes an HTL/CTL candidate whose native range overlaps a B-cell block already selected for the higher-priority class (5O1R, 7RXP; see Discussion).

### S4. Re-execution on complete proteins.

Over the sequence of the four source proteins of the antigens without short T-cell blocks (excluding fHbp site 2, treated separately since it does not fit the length hypothesis):

1. CSP (8FDD, 6B5M, 7RXP). Crystallized fragments located within the NPNA/NVDP repeat region of full-length CSP (UniProt P02893): 8FDD 121-133, 6B5M 120-132, 7RXP 324-388

(re-derived by pairwise alignment against the full-length sequence, 90.8% identity over 65 residues, consistent with a different circulating strain). With the full-length protein as input, 12 T-helper and 22 T-cytotoxic candidates are accepted; the T-helper candidate (126-140) and the T-cytotoxic candidate (128-137) directly overlap the ranges of 8FDD and 6B5M, and 6 of the 12 T-helper and 3 of the 22 T-cytotoxic candidates additionally overlap the 7RXP range. The floor-effect hypothesis is confirmed in all three cases.

2. Nucleoprotein, site 2 (7STS). Crystallized fragment (111 amino acids) at native positions 49-173 of the full-length nucleoprotein (UniProt P0DTC9, 419 amino acids). Of the 56 T-helper and 43 T-cytotoxic candidates over the full-length protein, 18 and 10, respectively, overlap this range. The floor-effect hypothesis is confirmed.

3. gp41 (MPER, 4XAW). Native motif located by its conserved flank (NWF[N/D]ITNWLWYIK, HXB2 671-683) in full-length gp41 (UniProt P04578, residues 512-856 of Env). Of the 38 T-helper and 26 T-cytotoxic candidates accepted over the full-length protein, none overlap this range. The closest candidate (151-158) leaves a single residue uncovered. The floor-effect hypothesis is not confirmed, but the closest candidate (ELDKWASL, CTL) contains the ELDKWA motif, the exact epitope sequence of the neutralizing antibody 2F5 (originally characterized by Muster et al. (1993), who define ELDKWA as the motif recognized by 2F5; conserved in 72% of HIV-1 isolates), directed against the same immunodominant region of gp41 (cluster II, aa 646-682 HXB2; Dorn et al. 2000; Morozov et al. 2022). Six T-helper candidates (123-137, 124-138, 126-140, 127-141, 128-142, 140-154) fall within that same cluster: the predictor did not reproduce the exact 4E10 epitope used in the panel, but it did correctly localize gp41's immunodominant region as a whole.

4. NHBA (5O1R). Crystallized fragment (115 amino acids) at native positions 373-488 of full-length NHBA (UniProt Q7DD37, strain MC58, 488 amino acids, C-terminal β-barrel domain). Of the 60 T-helper and 59 T-cytotoxic candidates over the full-length protein, 21 and 15, respectively, overlap this range. This confirms the floor-effect hypothesis, consistent with the other three cases in this section.

### S5. Complete phase-by-phase pipeline output for a representative structure.

To illustrate the pipeline's real intermediate output across every phase, this section walks gp120 CD4bs (HIV-1, PDB 3NGB) through every phase that produces new content beyond what the main text already shows. Unlike fHbp site 1 (5O14), 3NGB is the only antigen in the 17-structure panel that generates real, non-trivial output in every phase, including Phase 6 (HIV bnAb/CATNAP), which by pipeline design activates only for HIV Envelope antigens.

Phase 1/1.5 (sanitization/structure extraction): the 349-residue antigen sequence and coordinates were extracted from chain A of PDB 3NGB via the ATMSEQ route (default mode); this derived sequence, not the theoretical SEQRES, is the coordinate system used throughout this section.

Phase 2/3 (B-cell antigenicity, score fusion): Fig. S1 shows the fused score (all four engines contribute at every position for this structure input) and the resulting six Phase 3 candidate regions. The real VRC01 contact epitope (27 residues, recomputed in this session directly from PDB coordinates; NeighborSearch-style, ≤4.0 Å heavy-atom distance, excluding hydrogens and HETATM, antigen chain A against the correct Fab pair identified by real contact count rather than alphabetical chain order, which does not coincide with the biological pairing in this structure) falls into several clusters plus isolated contact residues. Candidate 225-234 (native 362-371) captures 5 of these residues, candidate 312-324 (native 455-467) captures 11, candidate 286-294 (native 429-437) captures 1 (native 430), and candidate 277-285 (native 420-428) captures none.

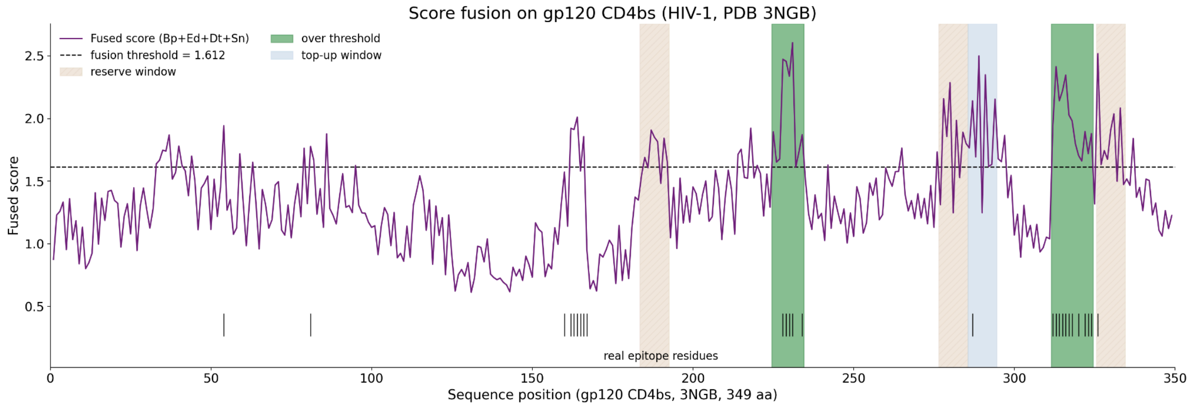


Fig. S1. Score fusion on gp120 CD4bs (HIV-1, PDB 3NGB). The fused score (sum of the four normalized engine scores) and the six Phase 3 candidate regions (shaded: over-threshold, top-up, or reserve windows; see Section 2) against the 27 real epitope residues (tick marks) computed by structural contact.

Phases 3b/4 (membrane topology and self-tolerance): Fig. S2: 318 of 349 residues (91%) resolve to TMbed's "o" (extracellular) class; 31 residues across 5 short segments are masked intracellular. One of the six Phase 3 candidates (312-324) partially overlaps a masked segment; since Phase 3b is informative only (Section 3.2 of the main body), this does not discard it. Four of the six Phase 3 candidates pass Phase 4 (70.0%, 66.7%, 0.0%, and 46.7% identity against the reference human proteome, all below the 75% discard threshold); the remaining two (77.8% and 88.9%) are discarded.

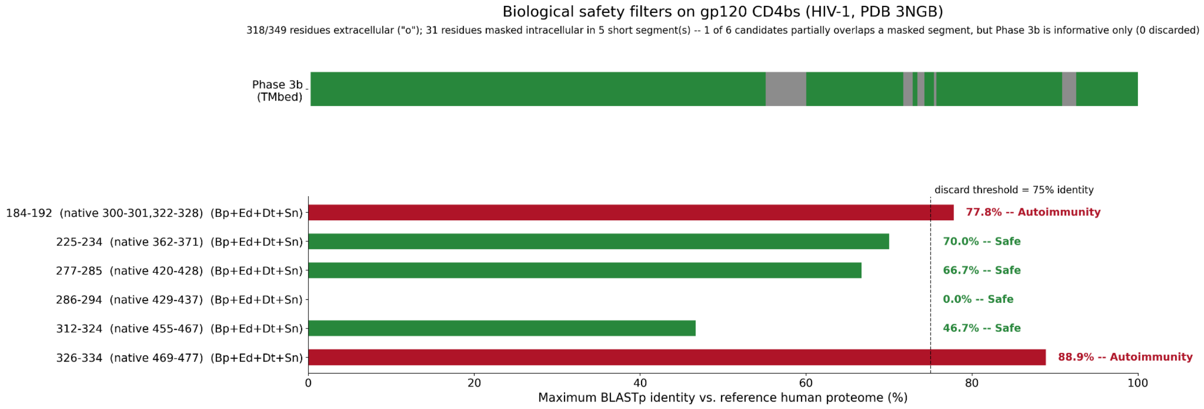


Fig. S2. Biological safety filters on gp120 CD4bs (HIV-1, PDB 3NGB). Top: TMbed topology track over the full 349-residue antigen (green = extracellular, gray = intracellular); 1 of the 6 Phase 3 candidates partially overlaps a masked segment, annotated but not discarded (Phase 3b is informative only). Bottom: maximum BLASTp identity against the reference human proteome for all 6 candidates, with the Phase 4 discard threshold (75%); 2 candidates (184-192, 326-334) exceed it and are discarded ("Autoimmunity").

Phases 4b/4c (allergenicity and N-glycosylation, informative only): Fig. S3: three of the four Phase 4 "Safe" candidates are flagged Allergen by AlgPred2's hybrid score (0.451, 0.363, 0.323), the fourth (277-285) is flagged Non-Allergen (0.229); one of the four contains canonical N-X-[S/T] sequons (312-324, 3 sequons, native positions N461, N462, N465).

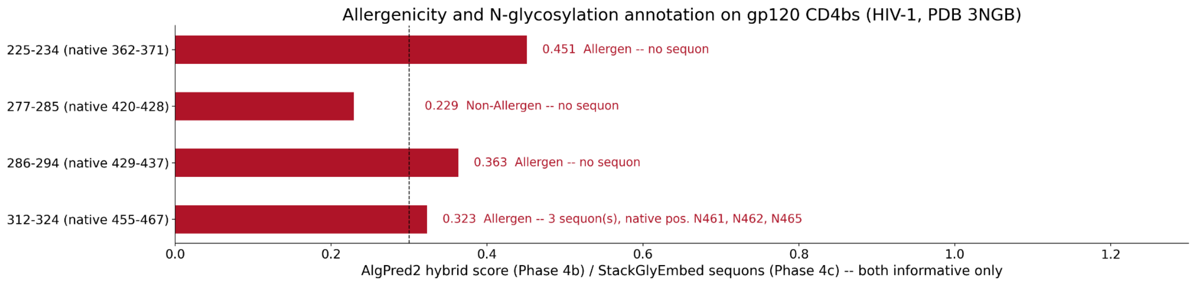


Fig. S3. Allergenicity (Phase 4b, AlgPred2 hybrid score) and N-glycosylation (Phase 4c, StackGlyEmbed) annotation for the 4 candidates surviving Phase 4 ("Safe") on gp120 CD4bs. Both phases are informative only and do not filter candidates.

Phases 5/5b (T-cell immunogenicity): already shown in Fig. 5 of the main text, which documents the 22 accepted T-helper and 37 accepted T-cytotoxic windows for this same structure, evaluated over the complete antigen independently of the B-cell candidates.

Phases 6/6b/6c (literature corroboration): Fig. S4. Phase 6 (HIV bnAb/CATNAP) applies, since 3NGB is an HIV-1 Envelope structure, and returns 5 specific hits (minimum 6-residue overlap) across two of the four candidates: 277-285 matches documented bnAb epitopes 2D3, JL413, G3-211, and G3-537; 286-294 matches 18A7. Phase 6b (cross-strain conservation) was not run for this structure (no variant panel supplied, an optional input, as for the rest of the panel; demonstrated separately for two of these candidates in Section S6, 8.0% and 0.2%). Phase 6c (IEDB) returns no specific hits for any candidate.

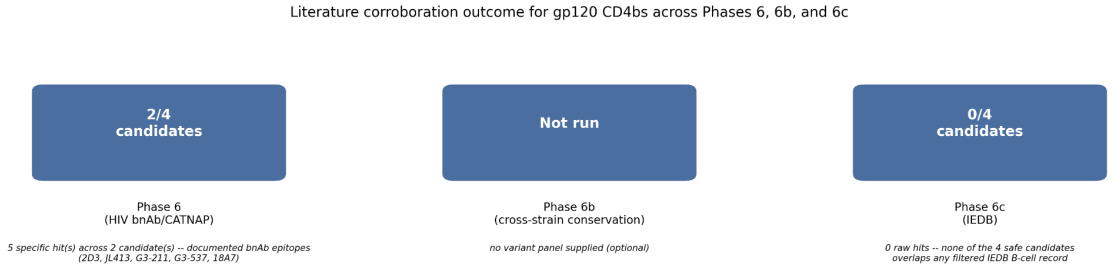


Fig. S4. Literature corroboration outcome for gp120 CD4bs across Phases 6, 6b, and 6c, evaluated over the 4 candidates surviving Phase 4. Phase 6 (HIV bnAb/CATNAP) finds 5 specific hits (≥6-residue overlap) across 2 of the 4 candidates, matching documented bnAb epitopes (2D3, JL413, G3-211, G3-537, 18A7); Phase 6b (cross-strain conservation) was not run for this structure (no variant panel supplied, optional); Phase 6c (IEDB) finds no specific hits.

Phases 7/8 (construct assembly and final check): Fig. S5: the assembled 155-aa construct (3 B-cell + 3 HTL + 3 CTL blocks + linkers, top-3 per class). The first B-cell block (source positions 309-327) recovers 12 of the 27 real epitope residues, the second (222-237) recovers 5 more, and the third (283-297) recovers 1 more, for a construct-level recall of 0.67, consistent with the retention figure reported in Section 3.3 of the main text. Phase 8's four additional checks on the fully assembled sequence flag it as Allergen, Non-Toxin, Moderate intrinsic antigenicity, and OTHER (no signal peptide); all four are informative only and do not filter the construct.

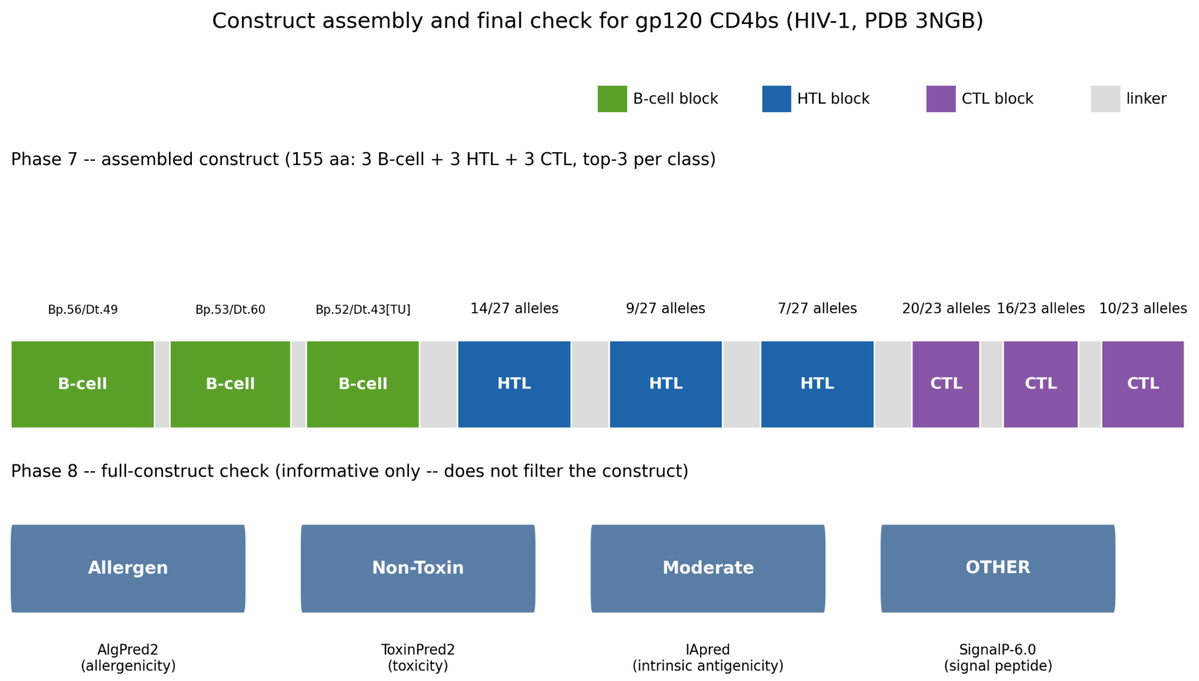


Fig. S5. Construct assembly and final check for gp120 CD4bs (HIV-1, PDB 3NGB). Top: the assembled 155-aa construct (3 B-cell + 3 HTL + 3 CTL blocks + linkers, top-3 per class). Bottom: Phase 8's four additional checks on the fully assembled sequence; all are informative only and do not filter the construct. B-cell block scores are BepiPred-3.0/DiscoTope-3.0 (Bp/Dt); [TU] = top-up window, [R] = reserve window (Section 2); blocks with no tag come from a region over the fusion threshold.

### S6. Cross-strain conservation against a real HIV Env panel.

Phase 6b (cross-strain conservation) was demonstrated for both HIV structures in the panel (3NGB, gp120 CD4bs; 4XAW, gp41 MPER) against a real, previously downloaded panel of 2,994 HIV-1 Env sequences from LANL/CATNAP (the same source already used for Phase 6), indexed locally with BLASTp (identity ≥90%, Section S1).

For 3NGB, two of its four Phase-4-surviving B-cell candidates show markedly different breadth: 225-234 (native 362-371) is conserved in 239 of 2,994 sequences (8.0%); 312-324 (native 455-467) in only 6 of 2,994 (0.2%), consistent with this second window's location in a more variable region of the CD4-binding-site loop. For 4XAW, the single B-cell candidate (residues 2-15 of the derived sequence) is conserved in only 7 of 2,994 sequences (0.2%) as evaluated.

This second figure is not a defect of the conservation methodology: Phase 6b now detects this case automatically. The candidate's BLASTp hits show a clean coverage cliff; near-total alignment coverage across residues 2-13, collapsing to under 1% over the last 2 residues; the signature of a non-native tail rather than genuine sequence variability (Section 2; verified panel-wide to avoid false positives on naturally hypervariable native candidates, see

Discussion). The pipeline flags the candidate, reports the supported sub-range (residues 2-13), and recomputes conservation over it alone: 1,799 of 2,994 sequences (60.1%), consistent with MPER's well-established status as a broadly conserved bnAb target. This is no longer a caveat requiring a manual check of the PDB deposition header; a check we found unreliable across the panel (the ENGINEERED/SYNTHETIC header flags fire on most structures regardless of whether the candidate is actually native, see Discussion), but an automatic, verified annotation (conservation_pct_native_core) alongside the raw figure.

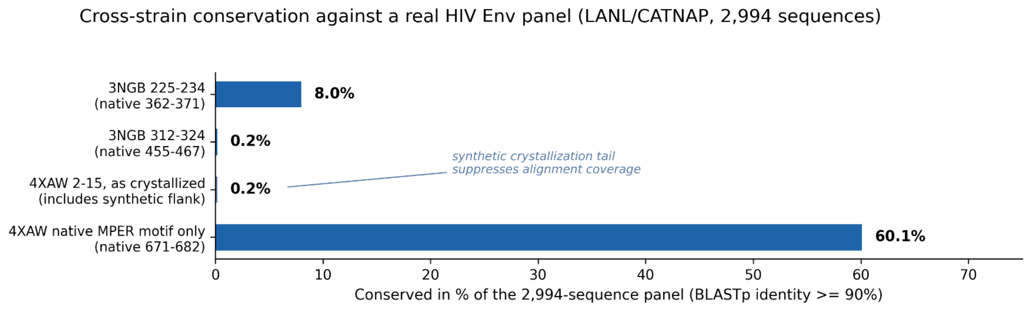


Fig. S6. Cross-strain conservation against a real HIV Env panel (LANL/CATNAP, 2,994 sequences) for the two HIV structures in the validation panel. 3NGB's two representative B-cell candidates (of its 4 Phase-4-surviving candidates) show markedly different breadth (8.0% vs 0.2%). 4XAW's single candidate, evaluated as crystallized (0.2%), is suppressed by a non-native flanking sequence added for crystallization; restricted to the native MPER motif alone, conservation rises to 60.1%, consistent with MPER's established status as a broadly conserved bnAb target (Section S6).

### S7. Leave-one-out cross-validation, fold by fold.

Table S7 reports the per-fold threshold and held-out performance of the 17-fold leave-one-out cross-validation described in Section 2, for the 4-engine score-fusion signal before the completion/reserve mechanism. The same threshold (1.612) was selected in 13 of 17 folds; the remaining 4 folds selected values within 3% of it (1.564-1.614). The pooled out-of-fold micro-F1 (0.297) is close to, and slightly below, the in-sample full-panel estimate at this fixed threshold (0.314), a modest, expected generalization gap at N=17. This pre-completion estimate is not directly comparable to the 0.335 micro-F1 reported in Table 2, which already includes the completion/reserve mechanism applied after threshold calibration.

| PDB | Fold threshold | Recall | Precision | F1 |
|---|---|---|---|---|
| 7CE2 | 1.612390 | 0.714 | 0.323 | 0.444 |
| 7OH1 | 1.604535 | 0.000 | 0.000 | 0.000 |
| 3NGB | 1.614260 | 0.407 | 0.846 | 0.550 |
| 4XAW | 1.612164 | 0.889 | 0.571 | 0.696 |

| | | | | |
|---|---|---|---|---|
| 5O1R | 1.612164 | 0.000 | 0.000 | 0.000 |
| 5O14 | 1.612164 | 0.000 | 0.000 | 0.000 |
| 8UP2 | 1.612164 | 0.000 | 0.000 | 0.000 |
| 2Q8A | 1.612164 | 0.524 | 0.478 | 0.500 |
| 2Q8B | 1.612164 | 0.571 | 0.333 | 0.421 |
| 9BJG | 1.612164 | 0.056 | 0.035 | 0.043 |
| 9BJH | 1.612164 | 0.000 | 0.000 | 0.000 |
| 8FDD | 1.612164 | 1.000 | 0.769 | 0.870 |
| 7RXP | 1.612164 | 0.067 | 0.067 | 0.067 |
| 6B5M | 1.612164 | 1.000 | 0.846 | 0.917 |
| 7STR | 1.564228 | 0.000 | 0.000 | 0.000 |
| 7STS | 1.612164 | 0.000 | 0.000 | 0.000 |
| 7BEP | 1.612164 | 0.423 | 0.204 | 0.275 |

Table S7. Leave-one-out cross-validation, fold by fold, for the 4-engine score-fusion threshold (Section 2), before the completion/reserve mechanism. For each of the 17 folds, we refit the threshold by maximizing micro-F1 over the exact grid of unique fused-score values on the other 16 antigens, then evaluated it, unmodified, on the held-out antigen (native PDB numbering; recall (TP/(TP+FN)), precision (TP/(TP+FP)), and F1 (their harmonic mean) computed per held-out antigen.

### S8. Structural validation panel: selection criteria.

Structures were selected purposively rather than through an exhaustive systematic search, prioritizing crystallographic structures where available (resolution 1.5-2.9 Å for 16 of 17 complexes; the sole exception, 7OH1, is a cryo-EM structure at 8.0 Å, noted as a caveat in Section 3.3). When more than one structure was available for the same antigen, we retained multiple entries when they targeted distinct epitope sites or antibody classes (the four AMA1 structures target three distinct antibodies; fHbp is represented by two distinct sites). We applied no pre-registered inclusion or exclusion protocol beyond structure availability and citation verifiability; the panel was designed for taxonomic and epitope diversity, not assembled through an exhaustive or randomized search of the PDB.

### S9. Computational requirements and runtime.

The full pipeline (8 phases, 14 external tools) ran on a 16-core, 11 GiB RAM machine without a GPU and required about 36 GB of disk space for all model weights and tool environments combined. A cold-cache, single-antigen run took 7m59s for the panel's shortest sequence (13 aa) and 7m37s for its longest (845 aa): wall-clock time in this range is dominated by per-tool startup overhead rather than sequence length. Running the full 17-structure panel end to end therefore takes on the order of two hours from a cold cache; subsequent re-runs on the same input reuse the content-hash cache described above and complete in under a second (Table S9).

| Resource | Measured value |
|---|---|
| CPU | 16 cores, no GPU (TMbed in fast mode) |
| RAM | 11 GiB on the validation machine |
| Disk (weights + environments, full install) | ~36 GB |
| Runtime, full pipeline, cold cache | 7m59s (13-aa antigen) to 7m37s (845-aa antigen) |

Table S9. Computational requirements and runtime, measured on the validation machine (16-core CPU, 11 GiB RAM, no GPU, TMbed run in fast mode). Runtime is a cold-cache, single-antigen, full 8-phase run (all 14 external tools), timed at the panel's two length extremes (13 and 845 aa); wall-clock time in this range was dominated by per-tool startup overhead rather than sequence length and was not separately measured for the remaining 15 antigens.

### S10. Score-fusion ablation excluding EpiDope.

Removing EpiDope and refitting the fusion threshold from scratch on the remaining three engines (BepiPred-3.0, DiscoTope-3.0, ScanNet) does not improve the fused signal: full-panel micro-F1 is comparable or slightly worse (0.300 versus 0.314), and the honest leave-one-out estimate is clearly worse (0.225 versus 0.297), driven by a large loss of precision (0.161 versus 0.287) that a higher recall (0.376 versus 0.308) does not offset. EpiDope is near-noise as a standalone engine, but its presence still shapes the fused signal's precision/recall balance in a way not recovered simply by excluding it and recalibrating; retaining it is an empirically neutral-to-favorable choice, not merely a reproducibility/licensing one.

### S11. Construct-level retention: score fusion versus BepiPred-3.0.

To test whether score fusion's residue-level advantage (Table 2) survives to the assembled construct, Phases 3-4-7 were repeated using BepiPred-3.0 alone (its own production threshold, the same completion/reserve mechanism, the same self-tolerance filter, and the same top-3 ranking). Construct-level retention is identical between the two methods (13/17), though they diverge on which two antigens they recover: fusion retains fHbp site 1 (5O14) but not AMA1 75B10 (9BJG); BepiPred-3.0 alone does the reverse (Table S11). Score fusion's residue-level precision/F1 gain does not by itself raise the construct-level retention count, but it does change which specific candidate region survives to the final ranking, case by case.

| PDB | Fusion TP/epitope | Fusion retained | BepiPred-3.0 alone TP/epitope | BepiPred-3.0 alone retained |
|---|---|---|---|---|
| 7CE2 | 11/14 | Yes | 14/14 | Yes |
| 7OH1 | 0/14 | No | 0/14 | No |
| 3NGB | 18/27 | Yes | 13/27 | Yes |
| 4XAW | 9/9 | Yes | 9/9 | Yes |
| 5O1R | 12/20 | Yes | 12/20 | Yes |
| 5O14 | 5/21 | Yes | 0/21 | No |
| 8UP2 | 0/12 | No | 0/12 | No |
| 2Q8A | 16/21 | Yes | 5/21 | Yes |
| 2Q8B | 14/21 | Yes | 17/21 | Yes |
| 9BJG | 0/18 | No | 4/18 | Yes |

| | | | | |
|---|---|---|---|---|
| 9BJH | 0/16 | No | 0/16 | No |
| 8FDD | 10/10 | Yes | 10/10 | Yes |
| 7RXP | 7/15 | Yes | 9/15 | Yes |
| 6B5M | 11/11 | Yes | 11/11 | Yes |
| 7STR | 2/16 | Yes | 8/16 | Yes |
| 7STS | 7/8 | Yes | 8/8 | Yes |
| 7BEP | 9/26 | Yes | 3/26 | Yes |

Table S11. Construct-level B-cell epitope retention (Phase 7's top-3 selection per class), comparing the production score-fusion pipeline against Phases 3-4-7 repeated with BepiPred-3.0 as the only antigenicity engine (same completion/reserve mechanism, self-tolerance filter, and top-3 ranking). TP, true positive residues (native PDB numbering) recovered in the final B-cell block(s); epitope, total real epitope residues for that structure. Retained, at least one true positive residue in the final construct. Both methods retain 13 of 17 structures, but not the same 13: fusion recovers fHbp site 1 (5O14) where BepiPred-3.0 alone does not, and BepiPred-3.0 alone recovers AMA1 75B10 (9BJG) where fusion does not.

### S12. Construct-level recall.

| PDB | Construct-level recall |
|---|---|
| 4XAW | 1.000 |
| 8FDD | 1.000 |
| 6B5M | 1.000 |
| 7STS | 0.875 |
| 7CE2 | 0.786 |

| | |
|---|---|
| 2Q8A | 0.762 |
| 3NGB | 0.667 |
| 2Q8B | 0.667 |
| 5O1R | 0.600 |
| 7RXP | 0.467 |
| 7BEP | 0.346 |
| 5O14 | 0.238 |
| 7STR | 0.125 |
| 7OH1 | 0.000 |
| 8UP2 | 0.000 |
| 9BJG | 0.000 |
| 9BJH | 0.000 |

Table S12. Construct-level B-cell recall (Phase 7's top-3 selection per class) by antigen, complementing the panel-wide figure (76.5%) reported in Section 3.3.
